# The tiniest genomes shrink much further: extreme reductive evolution in planthopper symbionts

**DOI:** 10.1101/2025.05.16.654412

**Authors:** Anna Michalik, Diego C. Franco, Junchen Deng, Monika Prus-Frankowska, Adam Stroiński, Piotr Łukasik

## Abstract

Strictly heritable endosymbiotic bacteria that provide limiting nutrients to sap-sucking hemipteran insects are known for their highly reduced genomes conserved in organization and function. Here, we show how in ancestral endosymbionts of planthoppers, *Sulcia* and *Vidania,* gradually losing genes during ∼263 my of co-diversification with hosts, co-infections by additional microbes and host ecological switches coincided with more dramatic genomic changes. At its extremes, this has resulted in the smallest non-organellar bacterial genomes known, at barely 50-52 kb. Such minuscule *Vidania* genomes evolved convergently in two planthopper superfamilies, and are strikingly similar in gene contents, including the ability to produce a single amino acid (phenylalanine) for the host. Losing many additional cell-function genes places them among mitochondria in the level of host dependence, further blurring the bacteria-organelle boundary.

**One-sentence summary:** We present the smallest bacterial genomes known and evaluate the patterns and processes of their reductive evolution

## Introduction

The smallest artificial, carefully designed bacterial genome capable of axenic growth in a rich medium is only 531 kilobases [kb], an order of magnitude less than that of the familiar *Escherichia coli* (Hutchison et al. 2016). However, for some two decades, we have known that the genomes of obligatorily intracellular insect-symbiotic bacteria that rely on host support mechanisms can be much smaller (Nakabachi et al. 2006). Specifically, the genomes of ancient, strictly heritable bacterial symbionts in diverse lineages of sap-sucking hemipteran insects have been convergently reduced to only encode genes involved in essential nutrient biosynthesis and some key cellular functions, sometimes barely exceeding 100 kb (McCutcheon and Moran 2012; Moran and Bennett 2014; Campbell et al. 2017; McCutcheon et al. 2024). (Bennett and Moran 2013, 2015; McCutcheon et al. 2019, 2024). Nevertheless, after the presumed early period of extensive genomic degeneration, several symbiont lineages have reached the stage where the remaining genes are indispensable and their further loss slows to nearly a halt (McCutcheon et al. 2019). This is demonstrated by the conservation of gene sets and order among divergent strains of *Buchnera* or *Tremblaya*, which have co-diversified with their respective hemipteran hosts for well over 200 my (Husnik and McCutcheon 2016; Chong and Moran 2018). However, stability has its limits, and surveys encompassing a greater range of host lineages have revealed lost genes and occasional rearrangements (Chong and Moran 2018; Deng et al. 2023) or more dramatic processes such as the extreme and rapid degeneration of the symbiont *Hodgkinia* in some cicadas (Campbell et al. 2017; Łukasik et al. 2018). Unfortunately, taxonomically limited sampling within most insect clades has prevented us from identifying broader patterns and key drivers of further genomic evolution and reduction.

Here, we uncover the genomic evolutionary patterns in planthoppers (hemipteran infraorder Fulgoromorpha), a ∼263 my old clade ancestrally associated with two strictly heritable bacterial lineages with highly reduced genomes – a Bacteroidetes bacterium *Candidatus* Sulcia muelleri (thereafter *Sulcia*) and a Betaproteobacterium *Candidatus* Vidania fulgoroidea (thereafter *Vidania)* (Urban and Cryan 2012; Deng et al. 2023, 2024). Using metagenomic data for 149 planthopper species representing 19 families, we ask about the processes, limits, and drivers of genomic evolution of these ancient symbionts.

## Results

### *Sulcia* and *Vidania* have co-diversified with planthoppers for ∼263 my

*Sulcia* and *Vidania* infections are broadly distributed across planthopper taxonomic diversity (Fig. 1). Based on rRNA sequences reconstructed from metagenomes, we have detected *Vidania*, usually accompanied by *Sulcia*, in 87 species of 149 sequenced, representing 15 of 19 surveyed families, on both sides of the deepest split in the planthopper phylogeny (Fig. 1, Table S1). Indeed, *Sulcia* infection is known to have pre-dated the divergence of planthoppers from their sister clade, Cicadomorpha, which includes cicadas, spittlebugs, leafhoppers, and treehoppers (Moran et al. 2005). In turn, the initial infection with *Vidania* seems to have occurred after these clades separated (Deng et al. 2023). *Sulcia* and *Vidania* have co-diversified strictly with their planthopper hosts (Fig. S1), as expected given their transovarial transmission (Michalik et al. 2009, 2021). Where both *Sulcia* and *Vidania* are missing, we always observe Hypocreales fungi – members of an insect-pathogenic clade known to have repeatedly replaced ancient bacterial nutritional endosymbionts of diverse Auchenorrhyncha, including planthoppers (Matsuura et al. 2018; Michalik et al. 2023; Siehl et al. 2024).

**Fig. 1.**
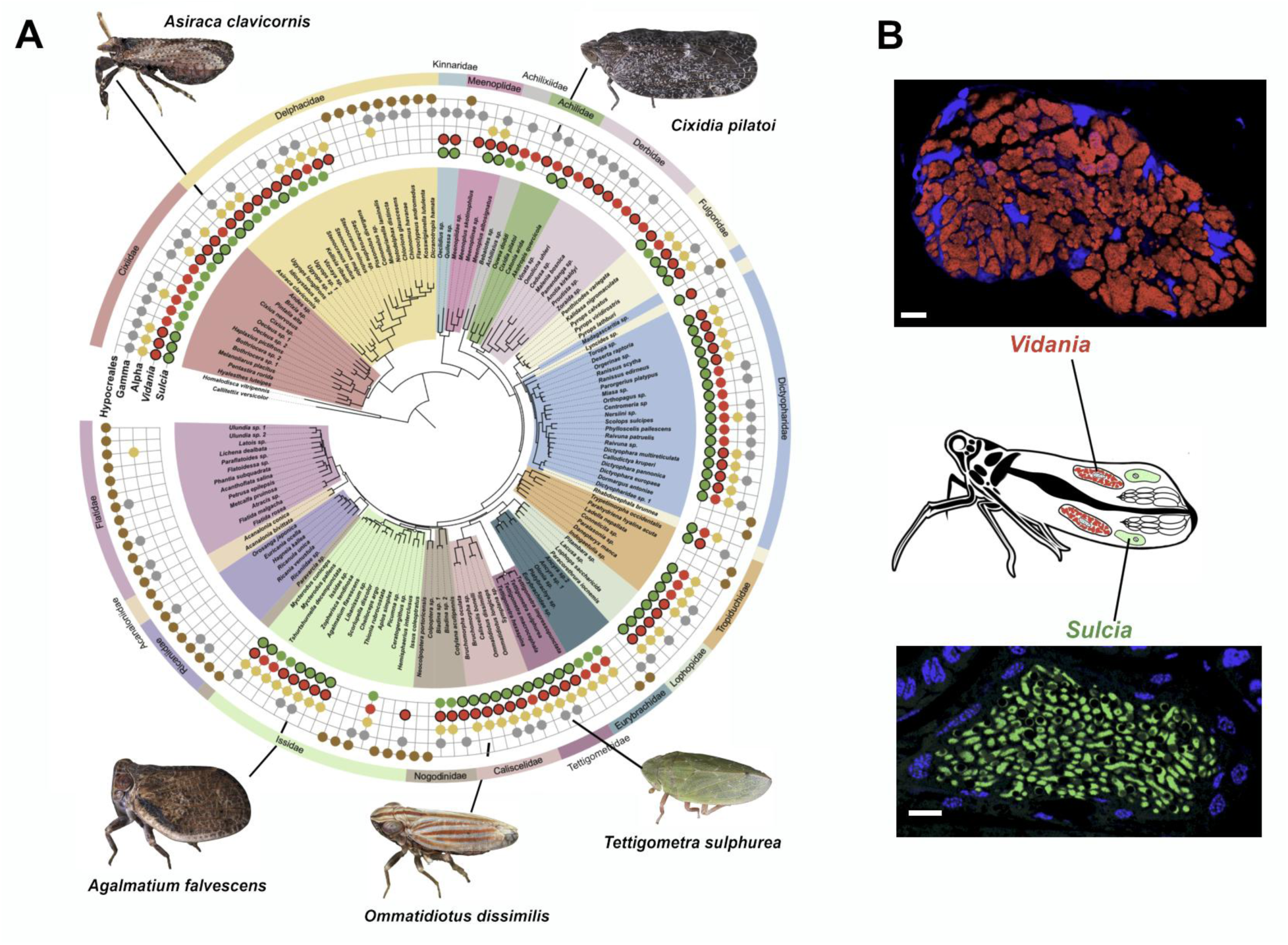
**A**. Two ancient symbionts, *Sulcia* and *Vidania*, are broadly distributed across planthopper taxonomic diversity. The planthopper phylogeny, based on 1164 nuclear and 13 mitochondrial genes, is based on (Deng et al. 2024). Red and green circles with bold outlines indicate strains whose genomes were fully assembled and used for subsequent analyses. **B.** *Sulcia* and *Vidania* live within the cytoplasm of dedicated cells (bacteriocytes) that form specialized organs (bacteriomes) that retain conserved organization across planthoppers, as visualized using confocal microscopy in *Akotropis quercicola* (Achilidae). Scale bar – 10 μm. Photo credit: G. Kunz (insects).

Further, we often find Gammaproteobacteria (*Sodalis*, *Arsenophonus*) and Alphaproteobacteria (*Wolbachia*, *Rickettsia*, Bartonellaceae, Acetobacteraceae), acquired independently by different host lineages (Michalik et al. 2021), known to provide vitamins and other complementary nutrients (Bennett and Mao 2018; Michalik et al. 2021, 2023, 2024) but also influence reproduction, defense, or other traits (Kaur et al. 2021; Davison et al. 2022). Together, our data show the relative stability of ancient bacterial associations while highlighting the evolutionary dynamics of more recently acquired microbes.

### *Sulcia* and *Vidania* have tiny genomes

Of 131 complete genomes of planthopper symbionts (64 *Sulcia*, 68 *Vidania*) that we used for content comparisons, most were circular. The two exceptions included the genome of *Sulcia* from *Trypetimorpha occidentalis*, previously confirmed to comprise two distinct circularly mapping contigs (Michalik et al. 2024), and *Sulcia* from *Brixia* sp., with the same organization. The analysis of read mapping and the remaining contigs in the assemblies assured us of the completeness of genomes, as did syntheny comparisons. Specifically, all but one of *Vidania* genomes, including the two smallest, were synthenic relative to the ancestral state (Deng et al. 2023) (Fig. S2). In *Sulcia*, one or more rearrangements relative to the ancestral state were seen in 13/63 (20.6%) genomes representing six families (Fig. S3).

The reconstructed *Sulcia* genomes ranged in size from 137,729 bp to 180,379 bp, and *Vidania* ranged from 50,141 bp to 136,554 bp (Table S2). Several newly assembled *Vidania* genomes are much smaller than the tiniest bacterial genomes characterized so far, including *Nasuia* from leafhoppers (≥107.8 kb), previously characterized *Vidania* (≥108.6 kb), or *Tremblaya* from mealybugs (≥139.0 kb) (Bennett and Moran 2013; Husnik and McCutcheon 2016; Vasquez and Bennett 2022; Michalik et al. 2024). Correspondingly, many *Vidania* strains encode fewer genes (as few as 63) than almost any other symbiont (Fig. 2A). Smaller gene sets have so far only been reported from *Hodgkinia* that form unique complexes of cytologically and genetically distinct but complementary lineages that apparently exchange gene products (Van Leuven et al. 2014; Campbell et al. 2017; Łukasik et al. 2018), and from organelles: mitochondria and some of the most reduced chloroplasts (Kelly 2021; Butenko et al. 2024). *Vidania* genomes were much more variable in size, gene set, and GC content than those of *Sulcia* (Fig. 2B).

**Fig. 2.**
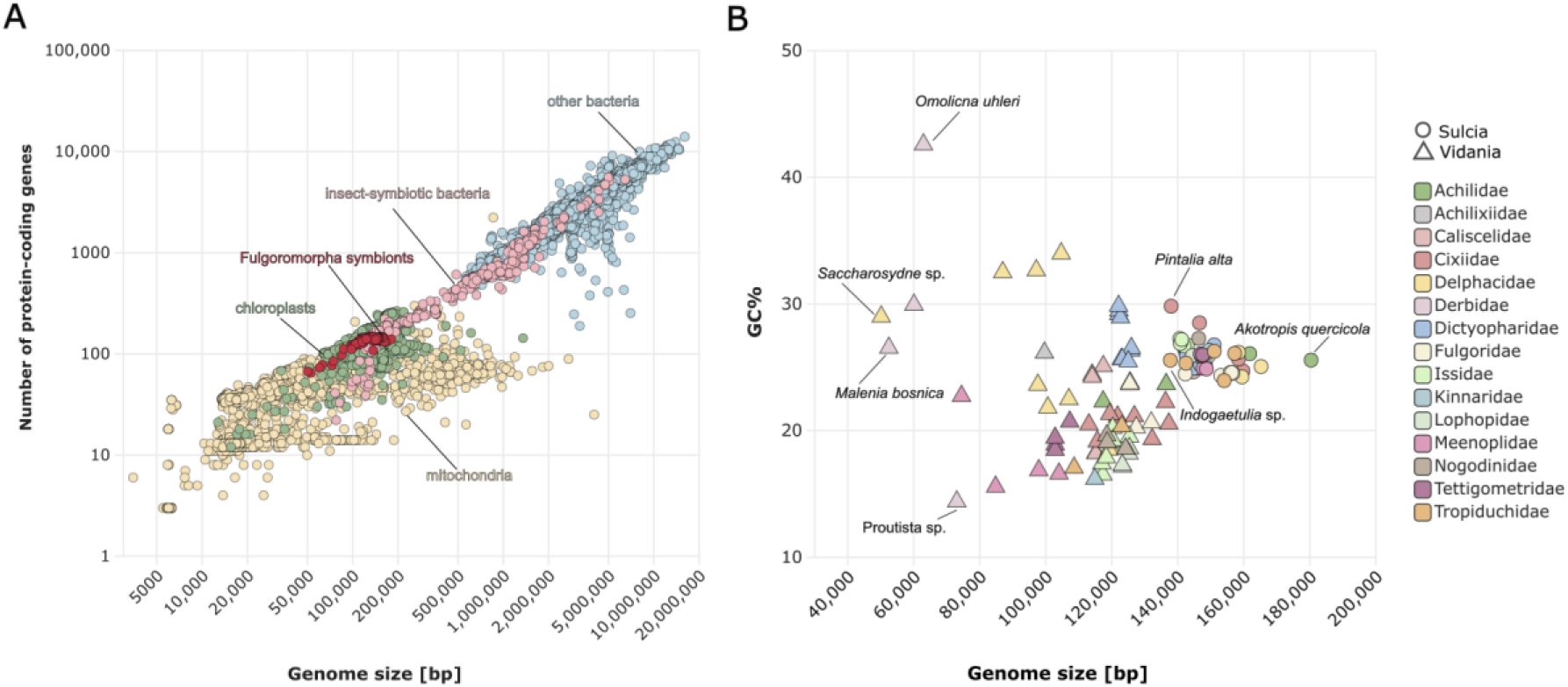
**A**. In terms of genome sizes and gene contents, planthopper symbionts (red) fall among the very smallest insect symbionts (pink), which in turn are among the smallest bacteria (blue), and among cellular organelles: mitochondria (orange) and chloroplasts (green). **B.** In terms of genome size and GC contents, *Sulcia* genomes tend to be relatively consistent, whereas *Vidania* genomes represent a much broader range on both scales.

### Variable rate of symbiont gene loss during co-diversification with planthoppers

The contents comparison among these unusual and, in the case of *Vidania*, extremely rapidly evolving genomes (Supplementary Text) enabled the reconstruction of their ancestral gene sets. In the last common ancestor of extant planthoppers, *Sulcia* must have encoded ≥164 total genes with identified functions. This includes 9 genes involved in the biosynthesis of three essential amino acids (Leu, Ile, Val) plus one gene for Phe biosynthesis, 132 genes involved in genetic information processing, and 18 involved in metabolism. The ancestral *Vidania* must have encoded ≥169 total genes, including 43 genes for the biosynthesis of seven amino acids (Met, Arg, Thr, Trp, Phe, Lys, His), 113 genes for genetic information processing, and 13 metabolism-related (Figs S4-S5). Additionally, across all *Sulcia* and *Vidania* genomes, we identified 40 and 86 open reading frames with no significant similarity to any records within the Uniprot database. While some of them were conserved in groups of species, suggesting functionality (Tables S3-S4), we did not include them in subsequent analyses.

We found no cases of gene gain by any lineage. However, gene losses were common during the evolution of both *Sulcia* and *Vidania*, with a notably higher loss rate in the latter (Fig. 3). In both symbionts, the patterns suggest a gradual loss of genetic information processing and metabolism genes during host-symbiont co-diversification. Some genomes lost <5% of the reconstructed ancestral gene set, but we observe several instances of much more substantial losses involving also nutrient biosynthesis pathways (Fig. 3A-B; Figs. S4-S5). For *Vidania*, this includes 49 gene losses on the branch leading to *Sacharosydne* sp. (Delphacidae), 40 losses in the lineage leading to the sole characterized Achilixiidae, *Bebaiotes* sp., 40 in the common ancestor of Derbidae (42), 32 in the ancestor of *Meenoplus skotinophilus* (Meenoplidae), and 25 in the ancestor of *Cixidia pilatoi* (Achilidae). Strikingly, four of the five lineages where *Vidania* lost most genes (all listed above except *Bebaiotes*) have also lost the *Sulcia* symbiont entirely, with only one additional recorded case of *Sulcia*-but-not-*Vidania* loss through apparent fungal replacement in *Issus coleoptratus* (Issidae) lineage. In turn, the branch leading to *Trypetimorpha occidentalis* (Tropiduchidae) and the ancestral branch of Tettigometridae exemplify cases where both *Sulcia* and *Vidania* lost multiple genes (3A-B; Figs. S4-S5).

**Fig. 3.**
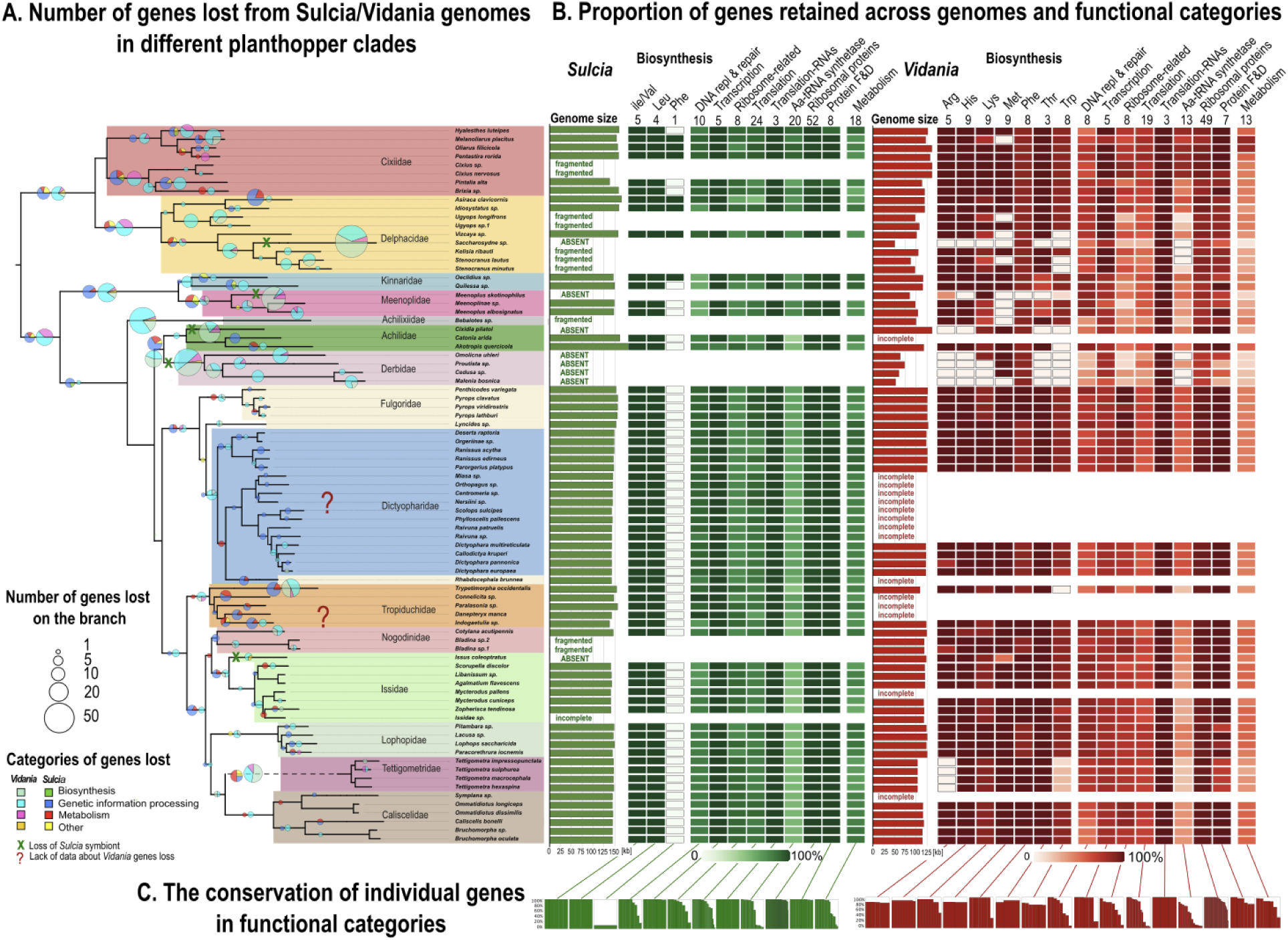
The reconstructed patterns of *Sulcia* and *Vidania* gene retention across the planthopper phylogeny indicate gradual gene losses during host-symbiont co-diversification, and much more dramatic changes in some lineages, with substantial variation among symbionts and gene categories. **A.** The insect phylogenomic tree, trimmed from Fig. 1 to only include species with a complete genome of at least one symbiont, is used to map the reconstructed numbers of gene losses, indicated by size-scaled pie charts for each symbiont on the branches. **B.** For all complete *Sulcia* and *Vidania* genomes and functional categories, we show the retained proportion of the ancestral gene set. **C.** The conservation of individual genes, i.e., the proportion of genomes where the gene is retained, varies among gene categories.

We also found major differences among gene functional categories in the overall level of conservation (Fig. 4C). In *Sulcia*, ancestral biosynthesis genes are always retained, with one exception: *aspC*, involved in phenylalanine biosynthesis, is found in *Sulcia* from seven species, whereas in most other planthoppers this gene is encoded within the host or *Vidania* genome instead (Fig. S4-S5). In *Vidania*, despite overall conservation, every biosynthetic pathway has been lost from at least one genome, with the Phe pathway most consistently retained. In both symbionts, rRNA genes are always retained, and ribosomal proteins together with genes involved in protein-folding and degradation are highly conserved. In contrast, aminoacyl-tRNA synthetases are the functional category most frequently lost.

**Fig. 4.**
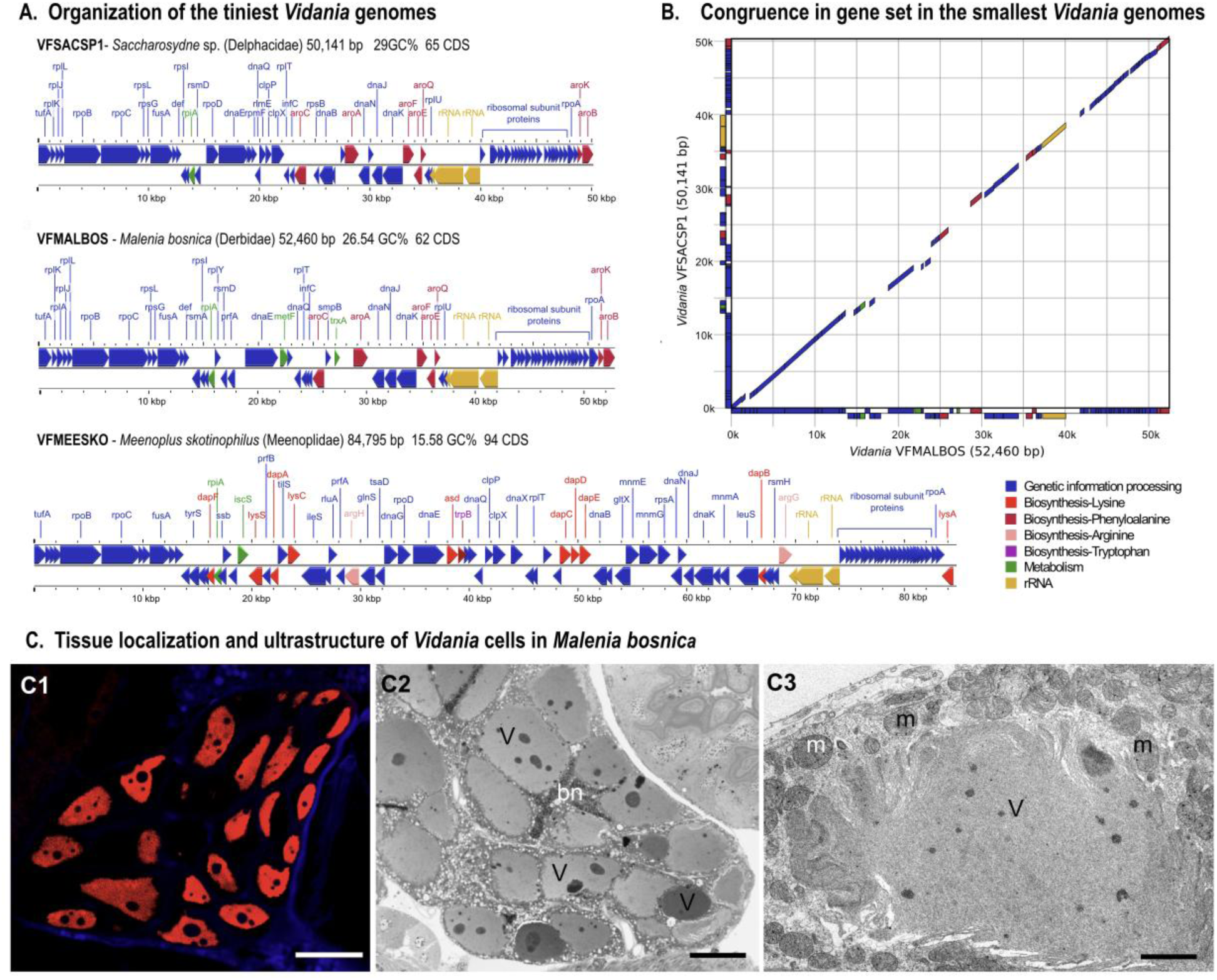
**A**. The representation of the smallest *Vidania* genomes from three planthopper superfamilies showcases their strongly reduced gene sets. **B.** The alignment of the two smallest *Vidania* genomes, which evolved independently from an ancestor approximately three times their size, shows striking convergence in their organization and content.. Due to the low sequence similarity among genomes, the alignment plot is based on the positions of annotated genes. **C.** Morphology and ultrastructure of *Malenia bosnica* bacteriome tissue and its tiny-genome *Vidania* VFMALBOS, as shown using fluorescence (C1), light (C2), and transmission electron microscopy (C3), differs from previously studied planthoppers by particularly high mitochondrial density. bn – bacteriocyte nucleus, m – mitochondrium, V – *Vidania* cell; C1, C2 – 10 um, C3 – 1 um.

### Convergence in the evolution of extremely reduced genomes

Planthopper symbionts include the smallest bacterial (non-organellar) genomes described to date: that of *Vidania* strain VFSACSP1 from *Saccharosydne* sp. (Dephacidae) has 50,141 bp, and that of VFMALBOS from *Malenia bosnica* (Derbidae) has 52,460 bp (Fig. 5A). VFSACSP1 encodes just 68 protein-coding genes with identified functions, and VFMALBOS 62. Notably, the extremely reduced state represented by these two genomes has evolved independently in two superfamilies separated by ∼263 my of evolution.

**Fig. 5.**
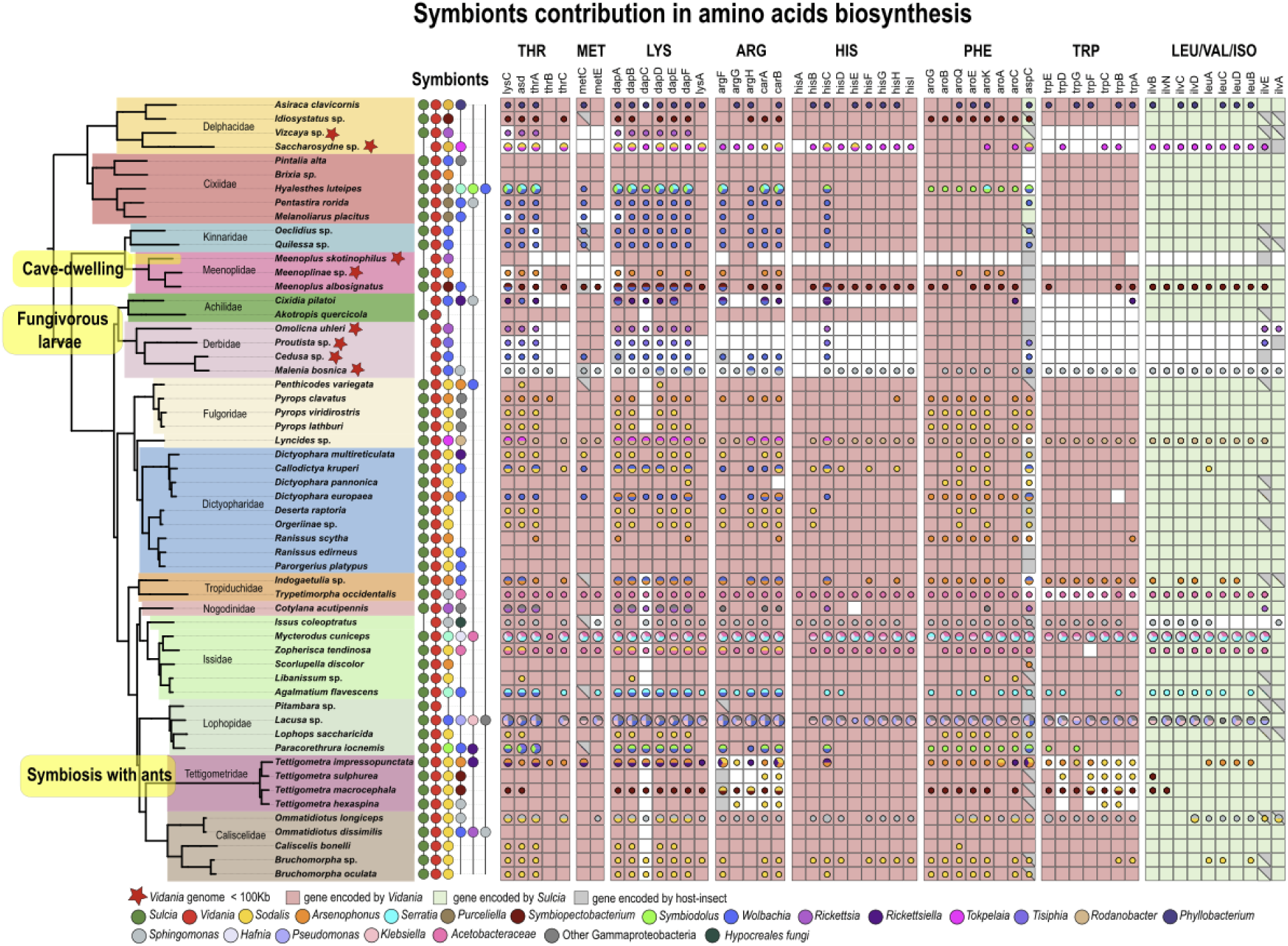
The distributions of amino acid biosynthesis genes across bacterial genomes in different planthopper species suggest compensation for the loss of pathways from *Vidania* genomes or the loss of *Sulcia*. Host phylogeny is redrawn from Fig. 1, but trimmed to only include species where *Vidania* and *Sulcia* (if present) genomes are complete. Bacterial presence and genus-level classification are based on the combined reconstruction of 16S rRNA and protein-coding genes. Green, red, and grey backgrounds in the main plot panel indicate genes present in *Sulcia*, *Vidania*, and the host genomes, respectively. Colored circles represent genes matched to other bacteria, with the circle size proportional to the number of genomes in which the gene was found.

Despite their independent origins, these genomes exhibit striking similarity in gene retention patterns and organization compared to their common ancestor, which must have had a genome approximately three times their size (Figs. 4A,B, S2, S5, S6). In both, the set of genes involved in essential cellular processes comprises rRNA genes, a set of 35-38 ribosomal proteins, 3-4 ribosomal DNA and RNA polymerase subunits, 15 other genes involved in genetic information processing, and one or two genes related to metabolism. Both these strains encode the entire ancestral set involved in phenylalanine biosynthesis, while pathways for the biosynthesis of six other essential amino acids have become entirely lost (Fig. 5B). Interestingly, neither genome retains any aminoacyl-tRNA synthetases. As in all other *Vidania* genomes, in VFMALBOS, but not in VFSACSP1, tRNAscan-SE identified some tRNA genes, although extreme rates of sequence evolution likely complicated their detection (Table S4). In contrast, the smallest genome from the third recognized planthopper superfamily, of *Vidania* VFMEESKO from a cave-dwelling *Meenoplus skotinophilus* (Meenoplidae), is substantially larger at 84,795 bp and retains an expanded set of genetic information processing and metabolism genes (Fig. 4A). It preserves the complete pathway for lysine biosynthesis and three genes from the arginine and tryptophan pathways, but interestingly, this is the only *Vidania* in our collection that has lost the phenylalanine biosynthesis pathway (Figs 4D, S6).

Notably, all planthoppers hosting these highly reduced *Vidania* strains have also lost *Sulcia:* they represent the few clades in our dataset where *Sulcia* was lost while *Vidania* remained (Figs 3, 5).

The organization and ultrastructure of *Vidania*-hosting bacteriome tissue of *M. bosnica* – the only tiny-genome species available for microscopy – differed from that of planthoppers studied before (Michalik et al. 2021, 2023, 2024) by exhibiting an unusually high density of mitochondria (Fig. 5C). Additionally, in some specimens we observed signs of cellular degeneration, including vacuolization of the bacteriocyte cytoplasm and higher bacterial cytoplasm density (Fig. 5C). In the bacteriome, we did not detect any other adjacent bacteria that could serve as an obvious source of proteins targetted to *Vidania* to replace the genes it lost (Bublitz et al. 2019).

### Nutritional functions explain the evolution of symbiont genomes and multi-partite symbioses

In hemipteran nutritional symbioses, the set of functions related to nutrient biosynthesis is generally conserved at the level of symbiosis, even when microbial partners change. In previously characterized planthopper symbioses, *Sulcia* and *Vidania* were thought to fulfill their hosts’ nutritional needs, jointly encoding most genes for the biosynthesis of all ten essential amino acids, with genes encoded on the host genome thought to complete the pathways (Fig. 5). However, we also find amino acid biosynthesis genes in other bacteria present in these metagenomes, including *Sodalis*, *Arsenophonus*, *Symbiopectobacterium*, *Wolbachia*, or *Tisiphia* (heritable endosymbionts previously detected in planthopper tissues (Michalik et al. 2023)), or *Serratia*, *Klebsiella*, *Sphingomonas*, or *Pseudomonas*, (likely less specialized gut associates). The genes they encode often overlap with those present in *Sulcia* or *Vidania*, but in many species that had lost *Sulcia* or some *Vidania* biosynthesis pathways, they often at least partially fill the resulting gaps (Fig. 5). This is most evident in two planthopper species with the tiniest *Vidania* genomes – *Saccharosydne sp.* and *M. bosnica* – where *Tokpelaia* and *Sphingomonas*, respectively, encode largely complete sets of biosynthesis genes. However, in assemblies for most other derbids, *Issus coleoptratus*, and others, we did not find alternatives for a large share of genes lost from ancient genomes. For example, for *M. skotinophilus*, none of the genes from the nine amino acid biosynthesis pathways lost were present in the 695Mb assembly for its dissected abdomen. We note, however, that functions encoded by microbes residing outside of tissue used for metagenomics, or sequenced to insufficient coverage, could have been avoided. Likewise, the reliable identification of biosynthesis genes encoded within the host genomes requires their more complete assembly.

The biology of most studied species is barely known, but mapping significant biological transitions onto the phylogeny also suggests the role of host ecology in structuring nutritional demands. Specifically, Derbidae and Achilidae feed on fungal hyphae during their immature stages (Howard et al. 2001), species in the genus *Tettigometra* engage in obligate trophobiotic associations with ants (Delabie 2001), and *Meenoplus skotinophilus* has adapted to life in caves (Hoch et al. 2025). The resulting alteration of nutrient availability and nutritional demands in these clades is likely to explain changes in biosynthetic gene sets.

## Discussion

The symbioses between planthoppers and their ancestral heritable endosymbionts *Sulcia* and *Vidania* can be remarkably stable, as shown by the similarity in symbiont genome organization and contents across the planthopper phylogeny and the previously demonstrated conserved organization of the symbiont-containing tissue (Michalik et al. 2023). These ancient symbionts have been gradually losing metabolism and genetic information processing genes during ∼263 my of co-diversification with hosts. However, the combination of contributions by other bacteria and of host ecological shifts can set the arena for a much more dramatic reduction of *Sulcia* and *Vidania* genomes and functions. In several planthopper lineages, these changes resulted in genomes much smaller than the tiniest characterized to date.

### How the tiniest genomes shrink further?

The mechanisms driving genomic reduction in insect nutritional intracellular endosymbionts include strong mutation and deletional pressure combined with the lack of recombination, no opportunities for gene acquisition, and strong genetic drift associated with symbiont transmission bottlenecks (McCutcheon and Moran 2012; Moran and Bennett 2014; McCutcheon et al. 2024). Through these processes, non-essential genes are gradually lost, with Mueller’s ratchet facilitating the fixation of somewhat detrimental but non-lethal changes (Moran 1996; Pettersson and Berg 2007; Bennett and Moran 2015). In ancient symbionts’ genomes, the long-term conservation of genes and pathways related to the biosynthesis of essential but limiting nutrients is linked to their essentiality for the hosts. However, such genes could be lost in three situations: if the host associates with additional microorganisms that provide the relevant function, host ecological changes render the function unnecessary, or the function is horizontally acquired by the host (Husnik et al. 2013; Husnik and McCutcheon 2016; Mao et al. 2018).

All three are known as significant evolutionary triggers, and our data suggest that at least the first two may drive symbiont genome degradation in planthoppers. Heritable endosymbionts co-infecting the same hemipteran species are often complementary in their biosynthetic capacity, including cases where genes lost by ancient symbionts are encoded by newer arrivals (Husnik and McCutcheon 2016; Bennett and Mao 2018; Michalik et al. 2024). Further, host nutritional shifts have been convincingly linked to microbiota changes – typically, obligate symbiont gains (Sudakaran et al. 2017; Cornwallis et al. 2023), but ancient symbiont losses from some Typhlocybinae leafhoppers that switched diets from plant sap to parenchymal cell cytoplasm (Buchner 1965) resemble our observations for Achilidae-Derbidae planthoppers.

Also, ancient heritable endosymbionts and organelles are known to rely heavily for basic cellular processes on proteins encoded within the host nuclear genome and targeted to organellar or symbiotic membranes or cytoplasm (Gray 2012; Smith and Keeling 2015; McCutcheon 2016, 2021; Mao et al. 2018; Kelly 2021; Husnik 2023; McCutcheon et al. 2024). It is hard to envision further losses of highly conserved symbiont metabolism or genetic information processing genes without host involvement, whether re-targetting products of genes previously encoded by the host, including those used to support mitochondria, or alternatively, the acquisition of new functions through gene duplication or horizontal gene transfer (Husnik et al. 2013; Duncan et al. 2014; McCutcheon 2021).

Unfortunately, we cannot unequivocally link these alternative mechanisms to specific genomic changes using the current short-read metagenomic data. For clades of interest, we would need dense sampling to comprehend the timing and order of host and symbiont genomic changes and the stability of other microbial associations, advanced tools such as spatial transcriptomics to understand cellular mechanisms, and improved understanding of species’ ecology.

### How small can bacterial genomes get?

To remain viable, a nutritional symbiont must provide biosynthetic functions that make it indispensable for the multi-partner symbiosis, and encode a set of genetic information processing and metabolism genes needed for its basic cellular functions that cannot be provided by the host (Moran and Bennett 2014; McCutcheon et al. 2024). We showed that in planthopper symbionts, nutritional functions encoded within a genome can be reduced to a single biosynthesis pathway, and the cellular function-related gene set to ca. 55 genes. Non-organellar genomes with more reduced coding capacity are only known from unique symbiotic complexes in cicadas, where genetically distinct cellular lineages derived from the ancestral *Hodgkinia* symbiont jointly encode the ancestral gene set and appear to exchange functions (Campbell et al. 2017; Łukasik et al. 2018).

The striking similarity among independently evolved tiniest *Vidania* genomes suggests that they may have reached the limits of reductive evolution in this specific system. However, judging from the parallels with mitochondria, symbiont genomes could potentially become much smaller. The most reduced known mitogenomes encode two rRNAs and two function-related proteins (Butenko et al. 2024), suggesting that with adequate host support, a theoretical gene content limit of three (rRNAs plus one function-related) may be enough to ensure the genome’s long-term persistence (Oborník and Lukeš 2015). Future genomic surveys across non-model insects may reveal symbionts closer to that limit.

### What are the consequences of extreme symbiont genome reduction?

Substantial gene losses can substantially change the symbiont’s role in insect biology. The biosynthetic capacity of the most reduced *Vidania* strains is limited to providing a single amino acid, phenylalanine. This precursor of tyrosine, needed for the formation, melanization, and sclerotization of a cuticle – insects’ first line of defense against natural enemies and abiotic challenges such as desiccation (Andersen 2010; Suderman et al. 2010; Kaltenpoth et al. 2025), has been identified as the key symbiotic contribution in beetles and hymenopterans (Vigneron et al. 2014; Anbutsu et al. 2017; Kiefer et al. 2023). We suspect that the ability to produce lysine, another important cuticle component, by *Vidania* from cave-dwelling *Meenoplus skotinophilus* may reflect host adaptation to an environment where a hardened, melanized cuticle is not necessary. Then, in these planthopper lineages that have also all lost *Sulcia*, the role of ancient nutritional symbiosis may have become reduced to supporting cuticle formation, whereas host insects’ other nutritional needs are fulfilled through altered diets or other microbes’ contributions.

Planthoppers’ reduced reliance on the ancient symbionts is likely to relax selective pressure on their genomic integrity and efficiency, driving ratchet-like accumulation of negative changes and functional losses. This will push the symbiosis deeper into what was described as “the evolutionary rabbit hole” – the state where the host is tied in a mutually obligate relationship with a degenerating partner (Bennett and Moran 2015). Our new results expand the range of potential long-term outcomes of such a situation. First, the symbiont undergoing reduction may reach a new ‘stable equilibrium’ at an even more reduced state, with some functions potentially shifted to other microbes or the host. The striking convergence and similarity among the two tiniest planthopper symbiont genomes suggest that such an equilibrium may exist for *Vidania*, and animal mitochondria demonstrate its potential of long-term stability (Kelly 2021). Second, the symbiosis may not be able to escape the degenerative spiral, losing functionality and efficiency, and perhaps ultimately leading to the extinction of the host lineage. The extremely fragmented and very rapidly evolving *Hodgkinia* symbiotic complexes of *Magicicada* periodical cicadas (Campbell et al. 2015, 2017), with their apparent high maintenance costs (Campbell et al. 2018), may exemplify this situation. Third, further host ecological changes may render the ancient obligate symbiosis unnecessary anymore: this is where some Typhlocybinae leafhoppers may have arrived (Buchner 1965). Finally, the replacement of degenerate ancient symbionts by newly arriving and more versatile and efficient bacteria or fungi (Matsuura et al. 2018; Michalik et al. 2023; Siehl et al. 2024) remains a possibility that may enable host organisms to take a new evolutionary path. On the other hand, such replacement is likely to initiate a new cycle of symbiosis degeneration (McCutcheon et al. 2019), as the “evolutionary rabbit hole” appears to be very difficult to escape permanently.

## Data availability

Raw sequencing data underlying this article can be accessed through NCBI Umbrella BioProject PRJNA684615, while the curated symbiont genomes are in the process of being uploaded. Genomes, genomic annotations, and protein references used for annotation were deposited in the Figshare repository under the link https://figshare.com/s/2184f9cfbaae8a21efff. Annotation, analysis, and visualization scripts are available through the project GitHub page: https://github.com/AnnaMichalik22/Planthopper-ancient-symbioses---supplementary-materials.git.

## Author contributions

**AM:** Conceptualization (equal), Resources (supporting), Methodology (equal), Investigation (lead), Formal Analysis (lead), Visualization (equal), Validation (equal), Data curation (equal), Funding Acquisition (equal), Project Administration (lead), Writing – Original Draft Preparation (supporting), Writing – Review & Editing (equal)

**DF:** Data curation (equal), Methodology (equal), Investigation (supporting), Formal Analysis (supporting),

**JD:** Investigation (supporting), Formal Analysis (supporting), Visualization (supporting), Data curation (equal)

**MP-F:** Methodology (supporting), Investigation (supporting),

**AS:** Resources (lead), Investigation (supporting),

**PŁ:** Conceptualization (equal), Methodology (equal), Software (equal), Investigation (supporting), Formal Analysis (supporting), Visualization (equal), Validation (equal), Funding Acquisition (equal), Project Administration (supporting), Writing – Original Draft Preparation (lead), Writing – Review & Editing (equal)

## Acknowledgments

We thank Chris Dietrich, Charles Bartlett, Hannelore Hoch, and Brian Fisher for providing specimens from their collections for this project, Arkadiy Garber for help with the data underlying Figure 2, and to Filip Husnik for valuable discussions. This study was supported by the Polish National Science Centre grants 2017/26/D/NZ8/00799 (to A.M.) and 2018/30/E/NZ8/00880 (to P.Ł.) and Polish National Agency for Academic Exchange grant PPN/PPO/2018/1/00015 (P.Ł.).

## Supplementary Appendix

**Figure S1.**
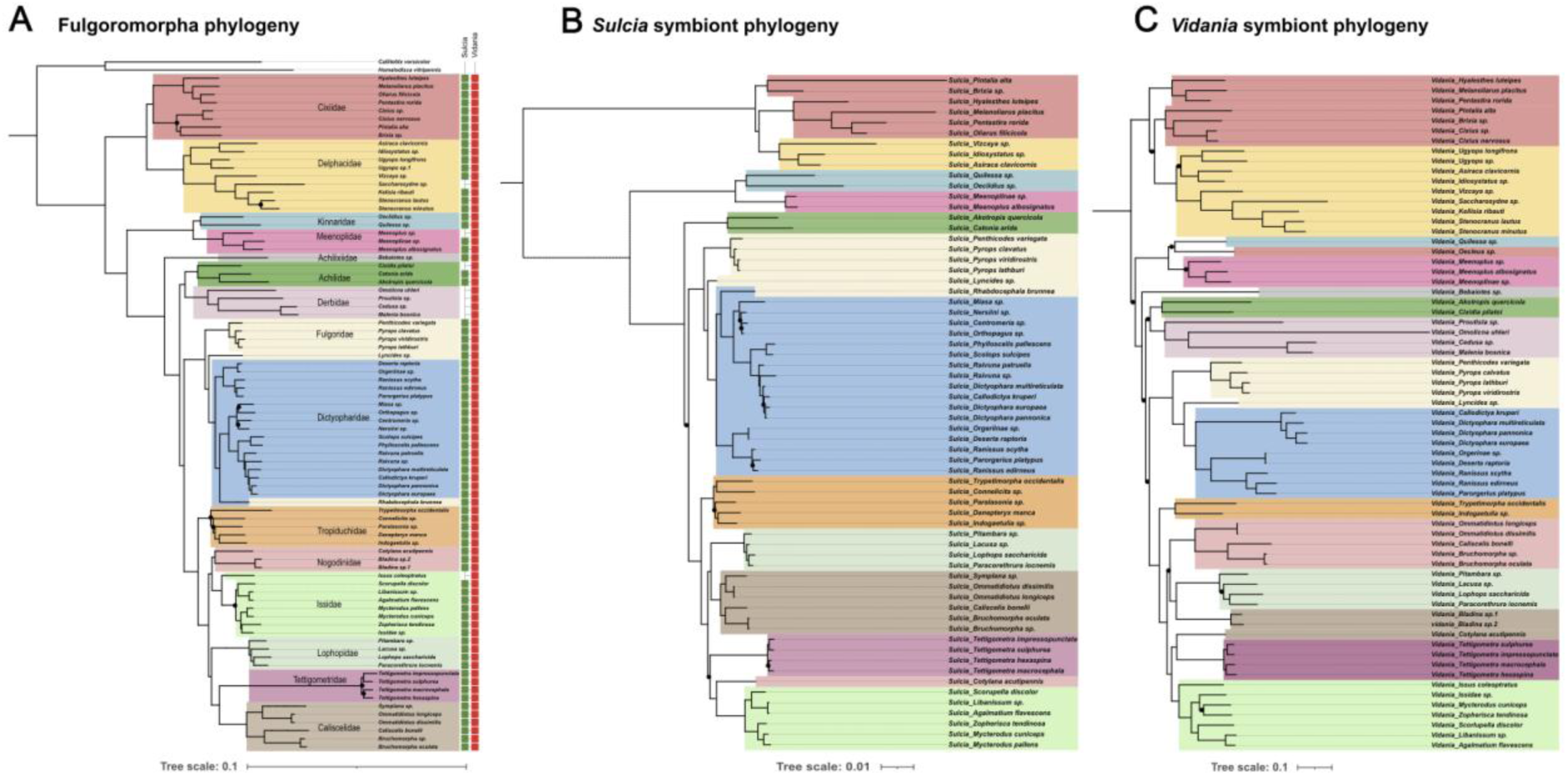
Co-diversification of planthoppers and their ancestral endosymbionts *Sulcia* and *Vidania*, based on genome-level data. The phylogenies of ancient bacterial endosymbionts *Sulcia* and *Vidania* essentially recapitulate that of their planthopper hosts, indicating strict co-diversification. **A.** The maximum likelihood phylogeny of 149 planthopper species based on 1164 nuclear and 13 mitochondrial genes (Deng et al. 2024); the topology is based on all species regardless of the infection state, but only the specimens that host either *Vidania* or *Sulcia* are shown; **B.** The ML phylogeny of 68 *Vidania* strains based on 98 genes; **C.** *Sulcia* ML phylogeny based on 130 genes. In all phylogenies, we used the same set of colors to indicate planthopper families. Black circles on the branches indicate bootstrap support values below 100.

**Figure S2.**
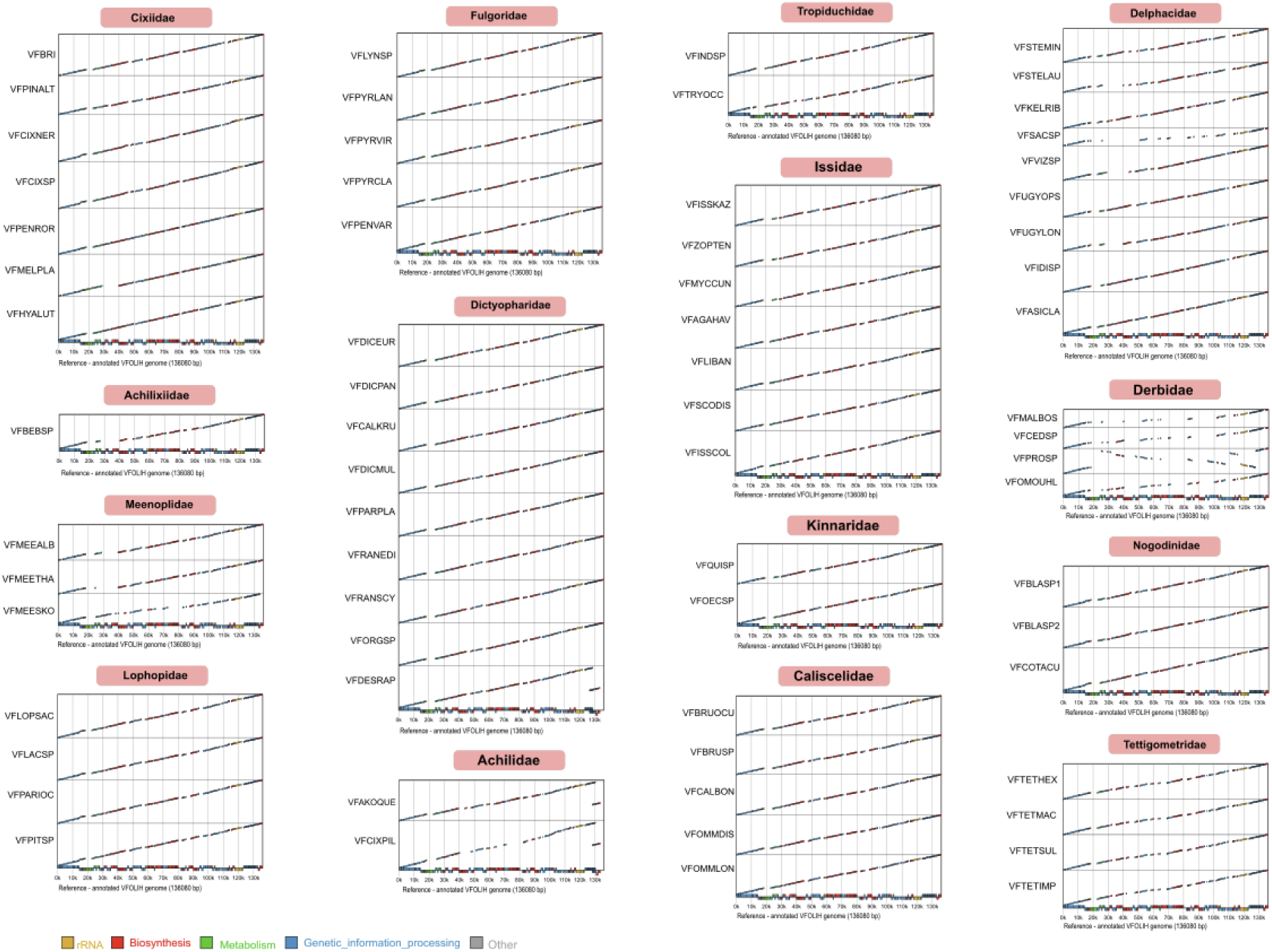
The alignment of all completed *Vidania* genomes against one of the most complete in the collection – that of *Vidania* VFOLIH from *Oliarus filicicola* (Cixiidae) – shows that the genomes generally retain conserved organization (with one exception – strain VFPROSP, Derbidae), but substantial portions of the genome are often lost. Alignment plots were made using a custom workflow that used annotation information.

**Figure S3.**
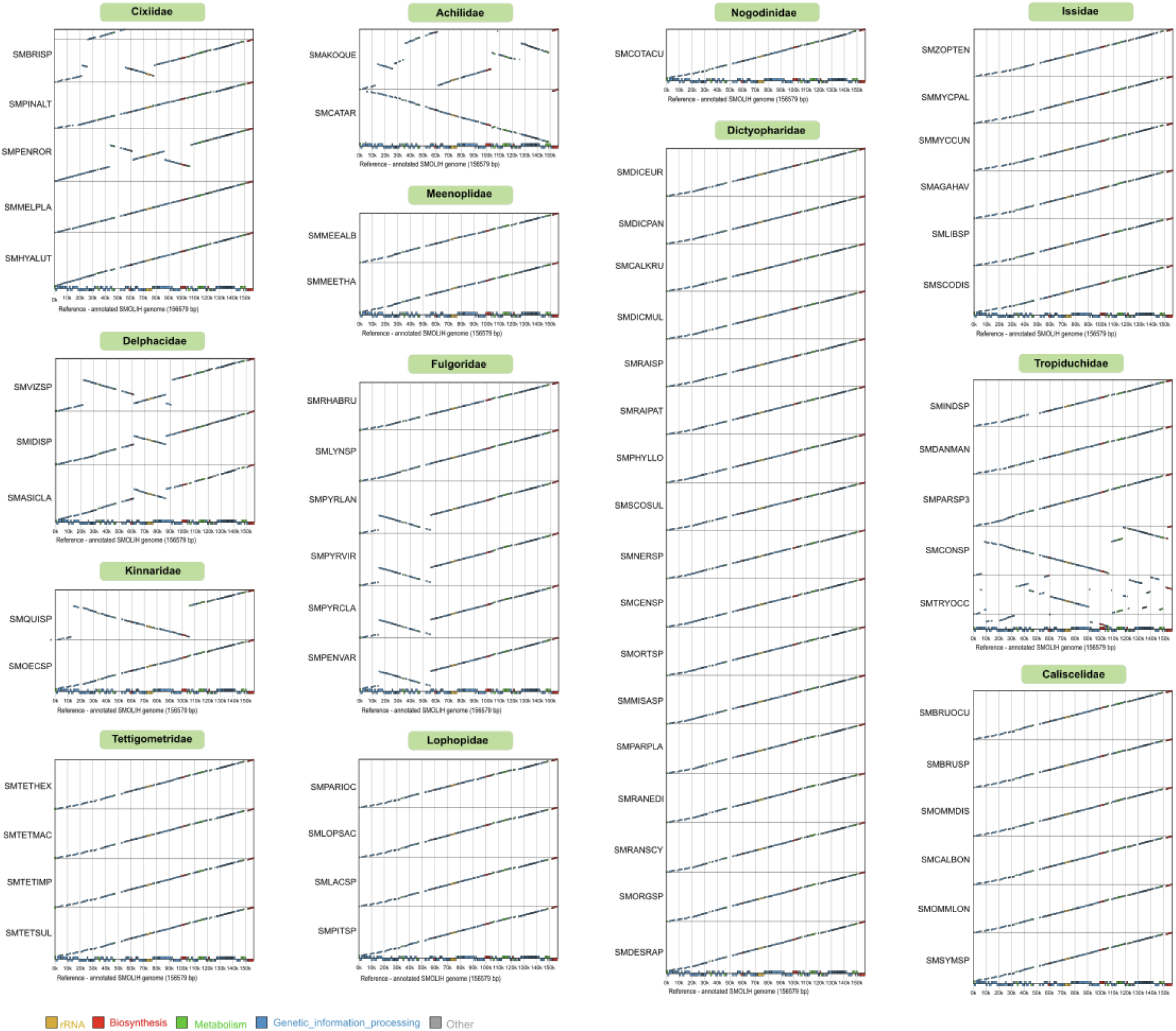
The alignment of all completed *Sulcia* genomes against one of the most complete in the collection – that of *Sulcia* SMOLIH from *Oliarus filicicola* (Cixiidae) – shows that the genomes usually retain conserved organization, but there have been multiple independent cases of genomic rearrangements, sometimes multiple. Alignment plots were made using a custom workflow that used annotation information.

**Figure S4.**
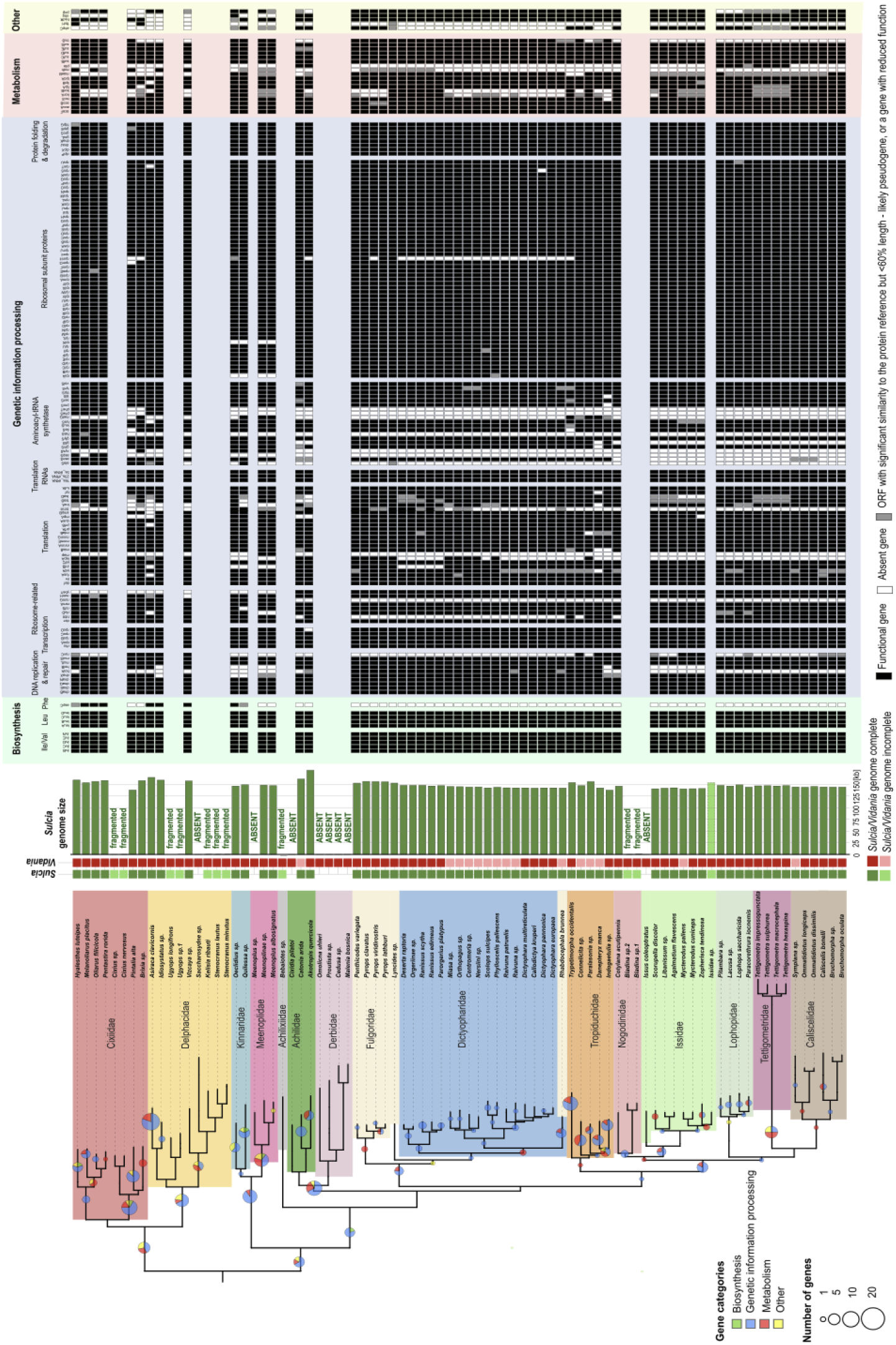
The distribution of genes across all complete symbiont *Sulcia* genomes from different planthopper species. As in Fig. 4, the insect phylogeny was redrawn from Fig. 1 but trimmed to retain only the clades where either *Sulcia* or *Vidania* genome is complete. For all complete *Sulcia* genomes and all genes with known functions, we show whether the gene was identified, and either deemed complete or truncated by >40% relative to the reference alignment (putative pseudogene). Circles on tree branches indicate the number of genes that were reconstructed as lost on a particular branch.

**Figure S5.**
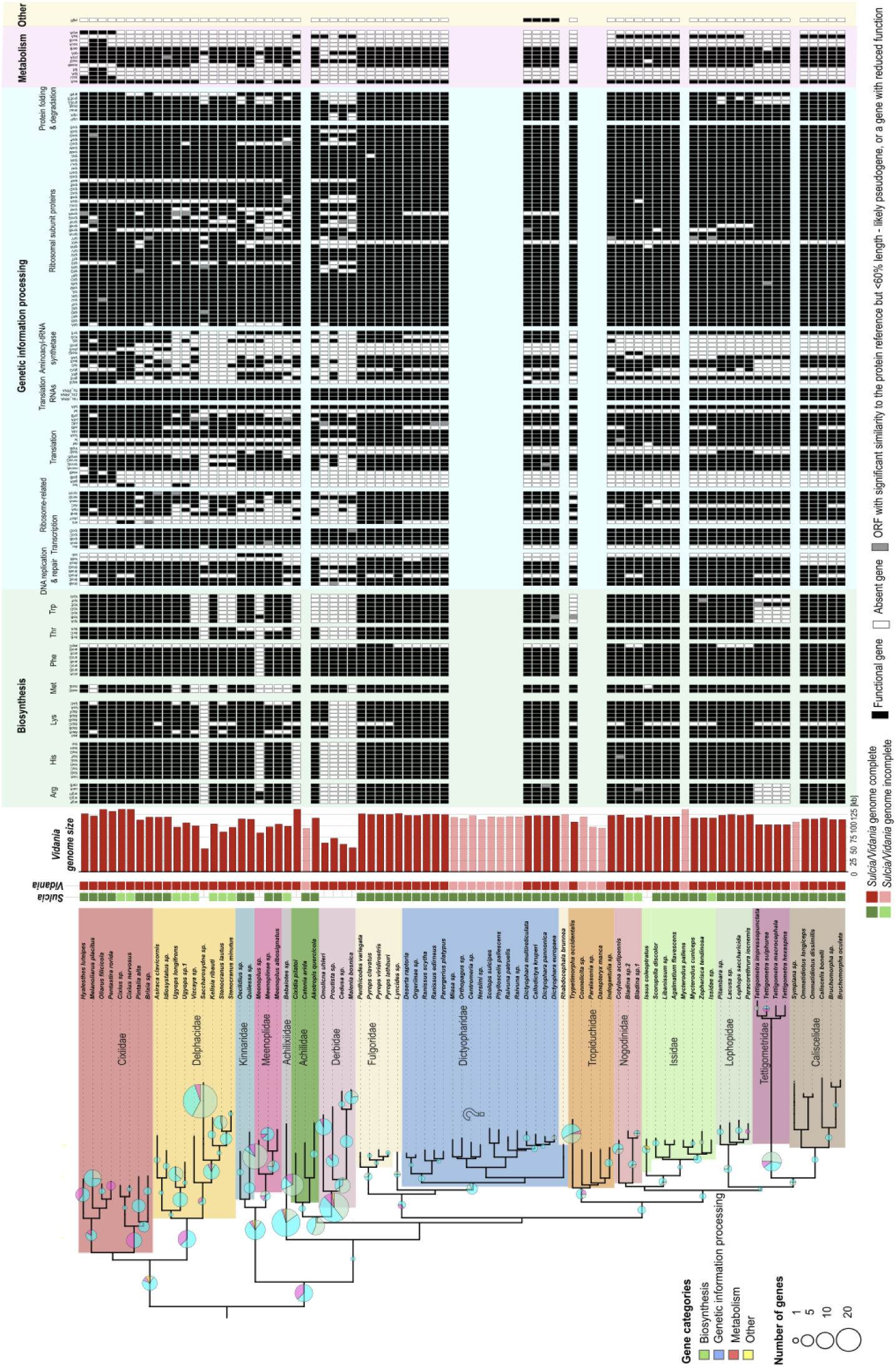
The distribution of genes across all complete symbiont *Vidania* genomes from different planthopper species. As in Fig. 4, the insect phylogeny was redrawn from Fig. 1 but trimmed to retain only the clades where either *Sulcia* or *Vidania* genome is complete. For all complete *Vidania* genomes, and all genes with known functions, we show whether the gene was identified, and either deemed complete, or truncated by >40% relative to the reference alignment (putative pseudogene). Circles on tree branches indicate the number of genes that were reconstructed as lost on a particular branch.

**Figure S6.**
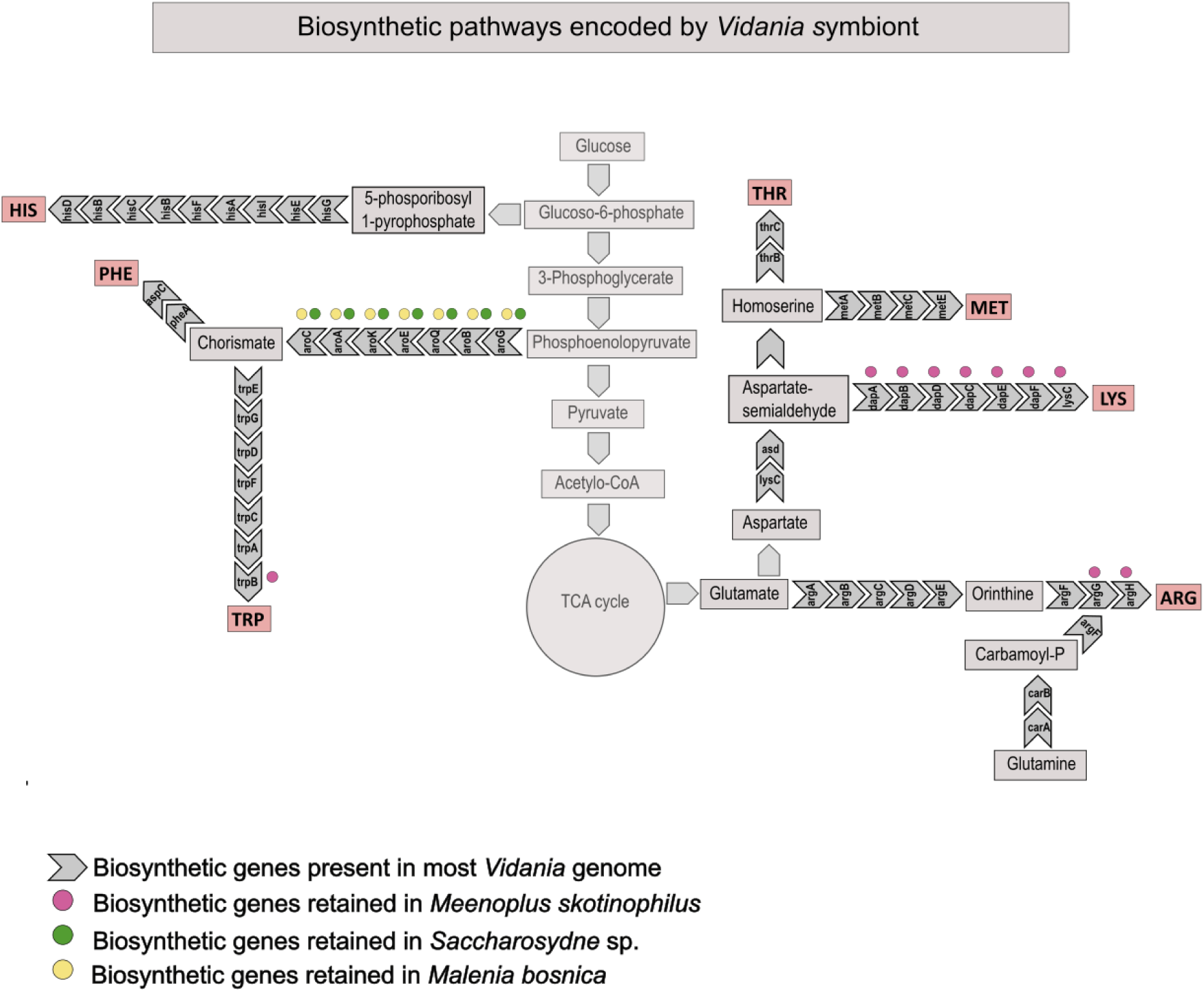
The representation of the ancestral amino acid biosynthetic pathways encoded by the ancestral *Vidania* – and the subsets encoded by the smallest *Vidania* genomes from three recognized superfamilies (VFSACSP1 from *Saccharosydne* sp., Delphacidae; VFMALBOS from *Malenia bosnica*, Derbidae; VFMEESKO from *Meenoplus skotinophilus*, Meenoplidae).

## Material and methods

### Insects

Individuals of 149 specimens representing 19 planthopper families were collected from their natural habitats around the globe between 2004-2021 (Table S1). Sampling was conducted in accordance with all applicable international and local permitting requirements. After collection, insects were identified based on morphological features and preserved whole in ethanol and stored at –20°C, or in some cases, partially dissected and fixed in a 2.5% glutaraldehyde solution and stored at 4 °C until further analyses.

### Metagenomic library preparation and sequencing

DNA from separated abdomens or dissected bacteriomes (symbionts containing-tissue) was extracted using the commercially available kits (Table S1) and used for metagenomic library preparation using the NEBNext Ultra II DNA Library Prep kit for Illumina (New England BioLabs) or Novogene NGS DNA Library Prep Set, with the target insert length of 150 bp. Due to the known issue of index swapping that occurs during cluster formation and sequencing on Illumina platforms (Illumina 2018; Costello et al. 2018) and which can lead to cross-contamination among samples in multiplexed lanes, we used dual-unique indexes for all but six earliest-processed samples as a means of reducing cross-talk. The library pools were sequenced in several batches on different Illumina platforms (Table S1), as previously described (Deng et al. 2024).

### Metagenome assemblies and characterization

Metagenomic raw reads were quality-filtered and adapter-trimmed using Trim Galore v0.6.4 (settings: –length 80 –q 30; https://github.com/FelixKrueger/TrimGalore). High-quality filtered reads were checked using FastQC v0.11.9 (https://github.com/s-andrews/FastQC). Contigs were assembled using Megahit v1.2.9 (maximum k-mer size = 255, min contig size= 1000) (Li et al. 2016). Contigs were validated, and their coverage was estimated by mapping reads against the assembled contigs. For the six libraries that were not double-uniquely indexed, we filtered the resulting assemblies for cross-contamination by discarding all contigs that had more than 10X (and typically >200X) greater coverage based on strictly mapped reads from another library with an overlapping index than based on reads from the same library.

Symbiont contigs were taxonomically identified through BLASTN and TBLASTX searches against the curated collection of *Sulcia* and *Vidania* genomes and the NCBI NT database. The circularity and contiguity of *Sulcia* and *Vidania* contigs were confirmed by read mapping and visualization using Tablet v. 1.20.12.24 (Milne et al. 2013), and based on the presence of overlapping ends. The circular *Sulcia* and *Vidania* genomes were rearranged so that they had the same orientation and start position (*Sulcia* – gene lipB; *Vidania* – tufA). Genomes of symbionts other than *Sulcia* and *Vidania* were represented by multiple contigs and will be presented and discussed in separate manuscripts.

### *Sulcia* and *Vidania* genome annotation

Because of the rapid evolution rate of *Sulcia* and especially *Vidania* symbionts and the extreme genetic distance from reference sequences, standard tools such as Prokka or Interproscan struggle to detect and consistently label genes, leaving annotation gaps and unannotated or hypothetical proteins. Therefore, *Sulcia* and *Vidania* genomes were annotated using a custom Python script modified from (Łukasik et al. 2018). The script extracts all Open Reading Frames (ORFs) and their amino acid sequences from each genome. It then searches these ORFs recursively using HMMER v3.3.1 (Eddy 2011) against custom databases containing manually curated sets of protein-coding, rRNA, and noncoding RNA (ncRNA) gene alignments from previously characterized *Sulcia* or *Vidania* lineages. We validated and updated these references after every round of annotation. rRNA and ncRNA genes were searched with nhmmer (HMMER V3.3.1) (Eddy 2011), and tRNAs were identified with tRNAscan-SE v2.0.7 (Wheeler and Eddy 2013). Based on the relative length compared to the reference genes, protein-coding genes were classified as putatively functional (≥60% of the reference alignment length) or truncated – likely pseudogenes (<60%). However, as explained in the section “Annotation challenges and unusual observations in fast-evolving symbiont genomes” below, the situation was not always clear-cut, and the decisions were made on a case-by-case basis.

Genomes for Figure 4 were visualized using Proksee (Grant et al. 2023). Comparative synteny plots were generated using a custom Python and Processing workflow modified from (Łukasik et al. 2018). Genome content tables were visualized using Processing and edited with Inkscape.

### Phylogenomic analyses

As the host phylogenomic framework, we used the planthopper phylogeny previously published by (Deng et al. 2024). The phylogenies of *Sulcia* and *Vidania* symbionts were reconstructed based on 131 and 99 protein-coding genes, respectively (Table S6). We chose *Sulcia* and the betaproteobacterial symbiont *Nasuia* from the treehopper *Entylia carinata* (ENCA) and the leafhopper *Macrosteles quadrilineatus* (ALF) as outgroups for *Sulcia* and *Vidania* phylogenies, respectively. The single-copy orthologs were identified using OrthoFinder v2.5.5 (Emms and Kelly 2019) based on the curated proteome resulting from the annotation pipeline above. Orthologs retained by >75% of samples (i.e., by >48 *Sulcia* lineages and by >53 *Vidania* lineages) were used for phylogenomic analyses. Nucleotide alignments of each ortholog were generated by MAFFT v7.526 (Katoh and Standley 2013). We kept only the 1st and 2nd codon positions in the alignments to avoid saturation effects from the 3rd codon position. Alignments were then concatenated and partitioned by genes. Phylogenies were inferred with the Maximum Likelihood (ML) approach in IQ-TREE2 v2.3.6 (Minh et al. 2020). To decide on the partitioning scheme and the substitution model, we asked IQ-TREE2 to perform extended model selection on each gene with free rate heterogeneity and subsequently merge genes until the model fit does not increase any further (setting: –m MFP+MERGE (Chernomor et al. 2016). The best partitioning scheme and the best-fit models were selected based on the highest BIC (Bayesian Information Criterion) scores. Bootstrapping was conducted using the approximate likelihood ratio test (SH-aLRT) and ultrafast bootstrap methods with 1000 replicates (Anisimova and Gascuel 2006; Hoang et al. 2018). Phylogenetic trees were visualised using iTOL (Letunic and Bork 2024).

### Analysis of symbiont contribution to amino acid biosynthesis pathways

The presence of bacterial amino acid biosynthesis genes in genomes of symbionts other than *Sulcia* and *Vidania* was assessed through HMMER searches within six-frame translated assemblies using HMM profiles from the NCBI Protein Family Models database. We taxonomically classified the identified proteins through BLAST-based comparisons against the nt database and against curated *Sulcia* and *Vidania* genomes. At that stage, we also filtered out genes of the known DNA extraction reagent-derived contaminant in our data, *Cellulosimicrobium*. The resulting hits were filtered based on alignment length and sequence similarity, with a large subset manually verified for gene identity and bacterial taxonomic assignment through blastn searches against the NCBI nt database. The results were visualised using a custom Processing script and edited in Inkscape.

### Microscopic analyses

#### Histological and ultrastructural analyses

The whole insect abdomens or dissected bacteriomes were fixed in 2.5% glutaraldehyde in 0.1 M phosphate buffer (pH 7.2) at 4℃. The fixed material was then rinsed three times in the same buffer with the addition of sucrose (5.8 g/100 ml) and postfixed in 1% osmium tetroxide for 2 hours at room temperature. After postfixation, samples were dehydrated in a graded series of ethanol (30-100%) and acetone, embedded in epoxy resin Epon 812 (Merck, Darmstadt, Germany), and cut into sections using Reichert-Jung ultracut E microtome. Semithin sections (1 µm thick) were stained in 1% methylene blue in 1% borax and analyzed and subsequently photographed under a Nikon Eclipse 80i light microscope (LM). Ultrathin sections (90 nm thick) were contrasted with uranyl acetate and lead citrate and examined and photographed at 80 kV using a Jeol JEM 2100 electron transmission microscope (TEM) at the Institute of Zoology and Biomedical Research, Faculty of Biology, Jagiellonian University.

#### Fluorescence *in situ* hybridization

Fluorescence *in situ* hybridization was performed using fluorochrome-labelled oligonucleotide probes targeting 16S rRNA of *Sulcia*: Sulc644 5’Cy3-CCmCACATTCCAGyTACTCC3’ (Koga et al. 2013) and *Vidania*: Bet940 5’Cy5-TTAATCCACATCATCCACCG3’ (Demanèche et al. 2008) symbionts. Ethanol-preserved insects were rehydrated and then postfixed in 4% paraformaldehyde for two hours at room temperature. Next, the material was dehydrated again by incubation in increased concentrations of ethanol (30-100%) and acetone, embedded in Technovit 8100 resin (Kulzer, Wehrheim, Germany), and cut into semithin sections (1 µm thick). The sections were then incubated overnight at room temperature in a hybridization buffer containing the specific sets of probes with a final concentration of 100 nM. After hybridization, the slides were washed in PBS three times, dried, covered with ProLong Gold Antifade Reagent (Life Technologies), and examined using a confocal laser scanning microscope Zeiss Axio Observer LSM 710 at the Institute of Zoology and Biomedical Research, Faculty of Biology, Jagiellonian University.

### Challenges and unusual observations in fast-evolving symbiont genomes annotation

The extreme evolutionary rates of ancient endosymbiont genomes have made their annotation challenging. The common challenges during the annotation of *Sulcia* and *Vidania* genomes included (1) marginal protein similarity between homologous genes; (2) the existence of phylogenetically conserved ORFs that corresponded to reference genes but were substantially truncated; (3) the merging of open reading frames (ORFs) from distinct genes into a single ORF, and (4) the presence of sequence variants that disrupted ORF of otherwise highly conserved genes. Below, we briefly summarize each of these challenges, present examples, and explain how we have considered such cases.

#### Very low similarity between homologous regions

In the rapidly evolving symbiont *Vidania*, sequences of the same protein from different strains sometimes had negligible similarity to each other. We established protein identity based on location within the largely syntenic symbiont genomes (Fig. S2-S3) and adjacent genes, on length, and on the result of phmmer searches against UniProt or SwissProt databases on https://www.ebi.ac.uk/Tools/hmmer/search/phmmer). However, when comparing such proteins, in many cases reciprocal blastp or phmmer, or hmmsearch with alignment of other *Vidania* gene copies as a reference, had low confidence scores or failed to identify matches at all. Low similarity also made it difficult or impossible to obtain acceptable protein sequence alignment for genes from different strains, and resulted in inconsistent annotations of such genes using hmmsearch searches with curated alignments for other copies of the same gene as a reference. In such cases, we created two or more variants of HMM references for the same gene, ensured that the ORF at the expected position is annotated when present and labeled consistently, and concluded that the successful detection of any of the variants indicates gene presence.

**Example:** The protein sequence similarity among variants of gene rplJ (50S ribosomal protein L10) in different *Vidania* genomes was often very low (<30% similarity in many cases). We established two alternative reference alignments, “rplJ” and “rplJ2”, verified that one of them is detected in all genomes where the ORF at the expected position is present, and concluded that the gene is present as long as one of the variants is detected.

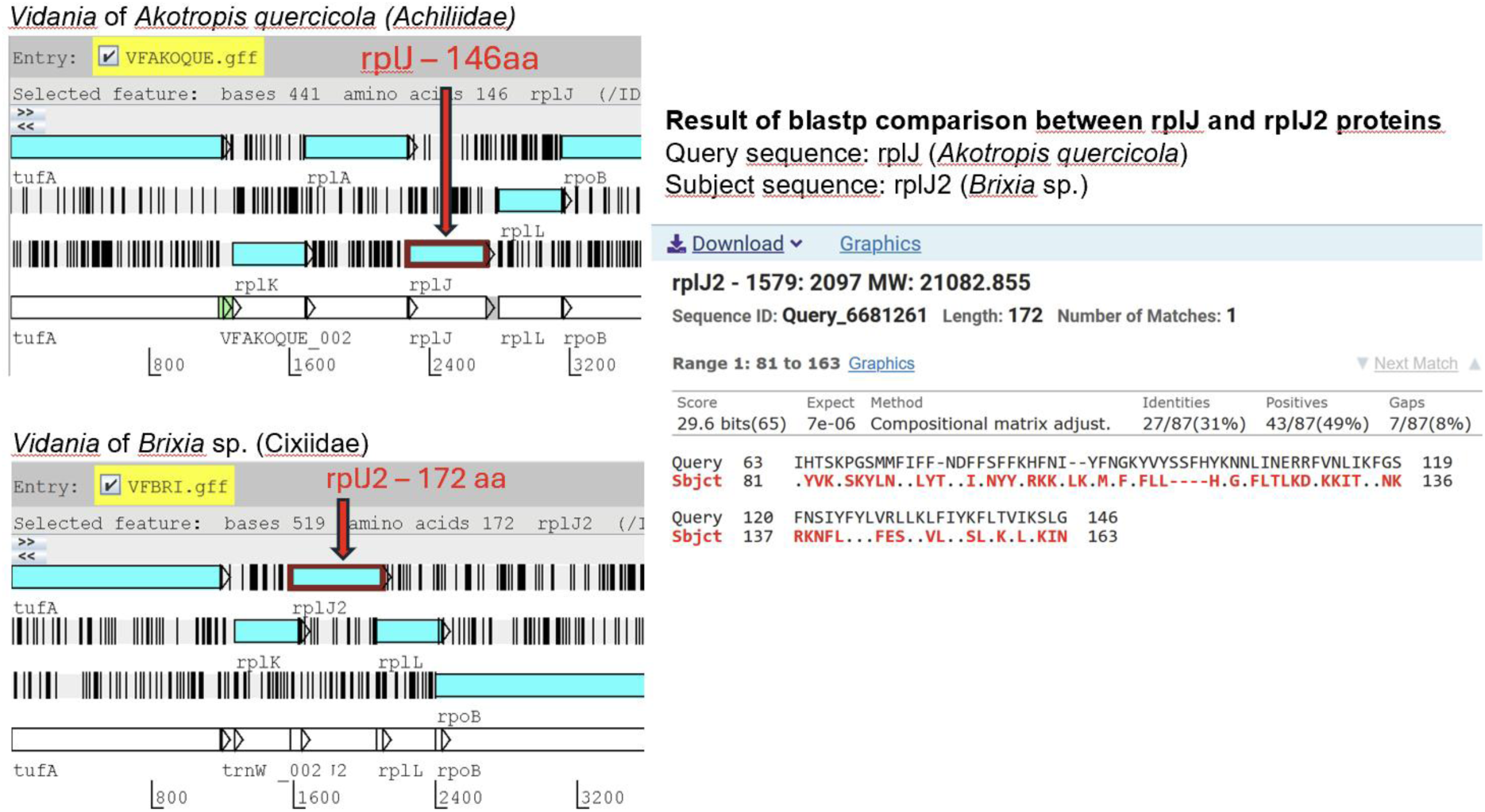

#### Gene truncation in *Sulcia* genomes

In *Sulcia*, certain genes were often substantially truncated relative to the most common length in the dataset, while retaining high sequence similarity. The truncation was often phylogenetically conserved, seen in all species from some clades, suggesting selective pressure for the retention of truncated variants and thus their functionality. We conclude that during the process of genome reduction, some gene domains may have been lost, but the truncated protein retained enough of the original function to justify their long-term retention. For such truncation conserved in at least 3 species, we established alternative reference alignments to ensure that they are scored as “functional”.

**Example:** The size of rplI gene (ribosomal protein) in *Sulcia* genomes from families Kinnaridae, Fulgoridae, Dictyopharidae, and Tropiduchidae is around 450 nucleotides, very similar to its length in *Eschericha coli* (447 nucleotides – https://www.uniprot.org/uniprotkb/P0A7R1/entry) whereas in the remaining nine families, we observed the truncation of this gene down to around 200 nucleotides.

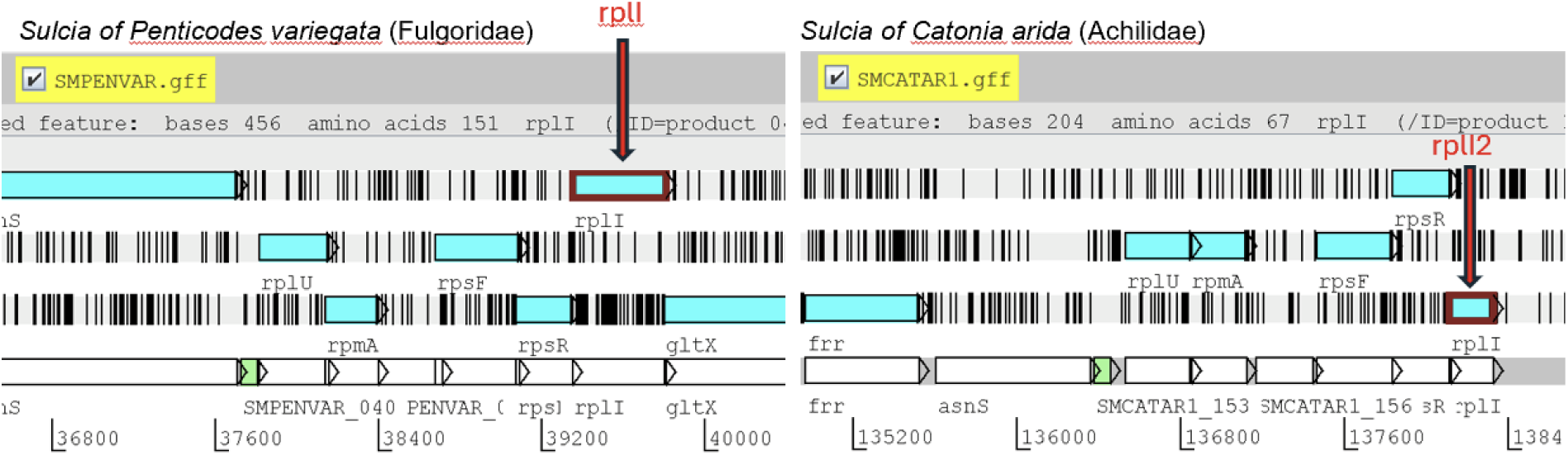

#### Open reading frames of distinct genes merged into one ORF

In *Vidania*, we observed cases when adjacent ORFs representing different genes, with very different functions, became merged into a single ORF. These cases were typically phylogenetically conserved – observed in symbionts in all species from a clade. Considering that the merged ORF contained nearly full-length coding sequences of the two genes, we concluded that both genes are likely functional and result in two distinct functional proteins. To ensure their successful detection, we created HMM references for the merged ORFs.

**Example:** In *Vidania* genomes from the clade comprising families Caliscelidae, Nogodinidae, Tettigometridae, Lophopidae, and Issidae, ORFs of genes tilS (tRNA(Ile)-lysidine synthase – a genetic information processing gene) and lysC (aspartate kinase – an amino acid biosynthesis gene), adjacent but separate in other planthoppers, became merged into a single large ORF. When such an ORF was detected, we scored both genes as present.

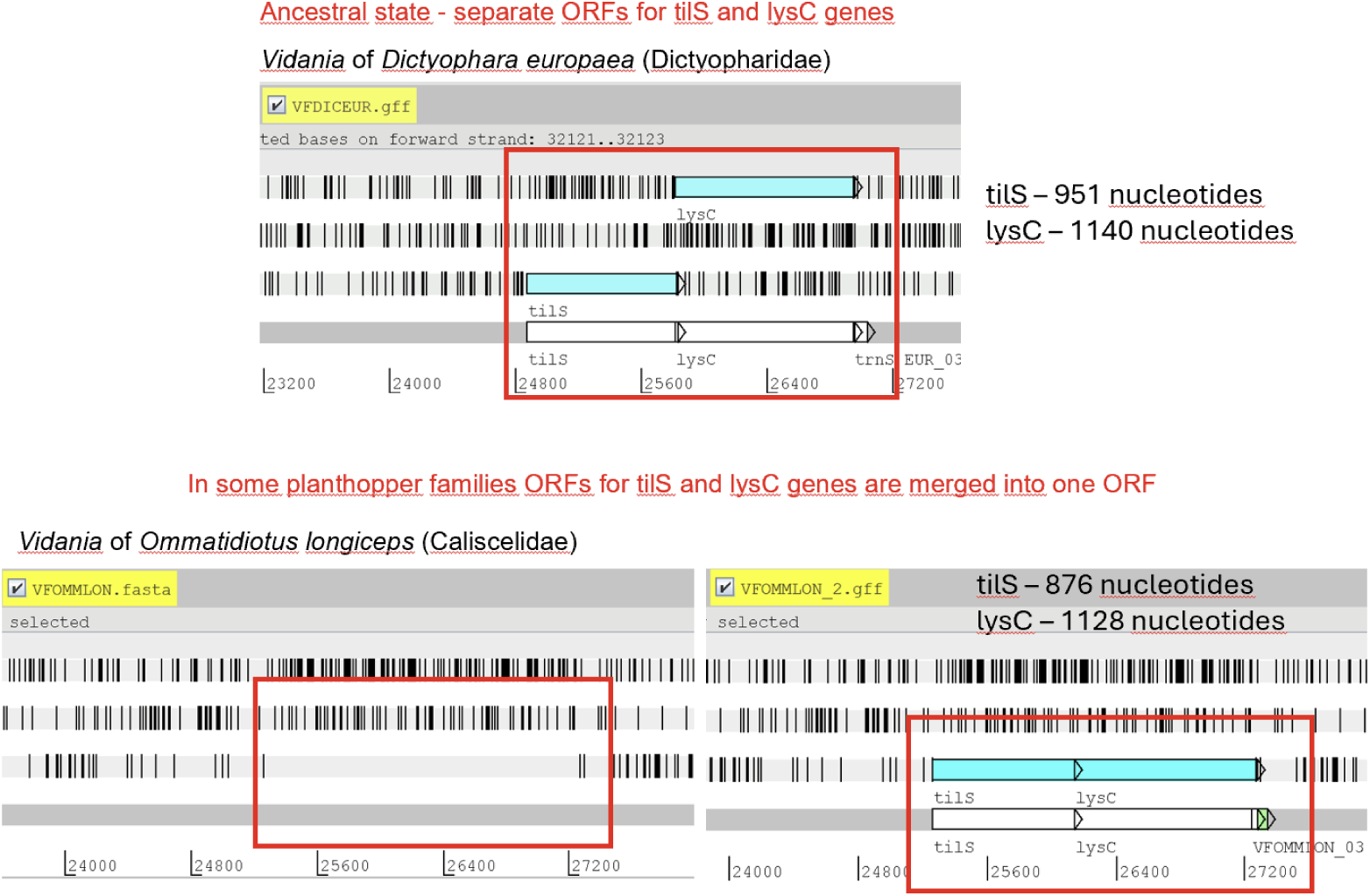

##### Other examples

– in *Vidania* genomes from the family Lophopidae, adjacent genes *dapC* and *dapD* were merged
– in *Vidania* genomes from the family Cixiidae, adjacent genes *maeB* and *aroE* were merged.

#### Open reading frames broken up into putative separate genes

In several genomes, we observed STOP codons or frameshifts within the open reading frames of conserved genes, confirmed as the dominant variant by inspecting mapped reads. In several cases, we excluded sequencing or assembly errors. We expect that in some cases, this might represent a recent mutation that has spread but not reached fixation within the highly polyploid symbiont genome, or within a population of symbiont cells. We also hypothesize that sequences may sometimes be corrected post-transcriptionally through an unknown mechanism. However, when considering such genes, we used the standard criteria, where ORFs of at least 60% of the reference length were scored as functional, and the shorter ones were scored as “truncated – putative pseudogenes”.

**Example:** Gene *dnaE* (DNA polymerase III alpha subunit) in *Vidania* of *Zopherisca tendinosa* is broken by an apparent frameshift into two separate ORFs, the longer of which was ca. 72% of the reference length. In this case, we classified the gene as functional.

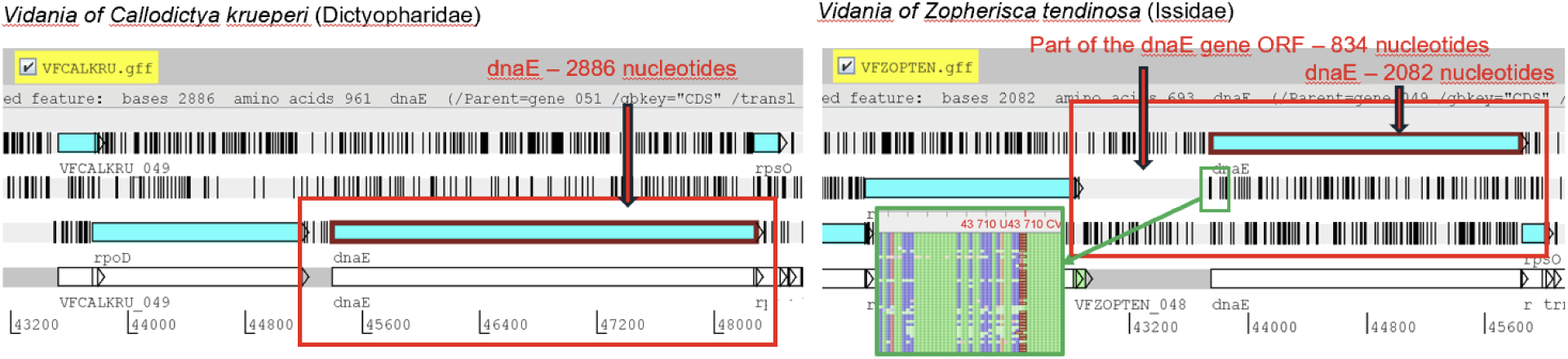

