## Supplementary Appendix for "The tiniest genomes shrink much further: extreme reductive evolution in planthopper symbionts"

**List of Supplementary Tables** 1

**Table S1.** List of planthopper specimens used in phylogenomic comparisons, their collection details, and microbial infections reconstructed using Phyloflash. 1

**Table S2.** The basic characteristics of 64 *Sulcia* and 68 *Vidania* genomes reconstructed from metagenomes 1

**Table S3.** *Sulcia* genome contents: the reconstructed presence of genes in *Sulcia* genomes deemed complete 1

**Table S4.** *Vidania* genome contents: the reconstructed presence of genes in *Vidania* genomes deemed complete 1

**Table S5.** Distribution of amino acid biosynthesis genes across symbionts in different planthopper species 1

**Table S6.** List of genes used for the reconstruction of *Sulcia* and *Vidania* phylogenies 1

The supplementary tables are available in the Figshare repository under the link https://figshare.com/s/2184f9cfbaae8a21efff

**Supplementary Figures** 3

**Figure S1.** Co-diversification of planthoppers and their ancestral endosymbionts *Sulcia* and *Vidania* 3

**Figure S2.** Gene content in complete *Sulcia* genomes across planthopper families 4

**Figure S3.** Gene content in complete *Vidania* genomes across planthopper families 5

**Figure S4.** Organization of *Sulcia* genomes in planthopper families 6

**Figure S5.** Organization of *Vidania* genomes in planthopper families 7

**Figure S6.** Amino acid biosynthetic pathways in reduced *Vidania* genomes 8

**Material and methods** 9

Insects 9

Metagenomic library preparation and sequencing 9

Metagenome assemblies and characterization 9
*Sulcia* and *Vidania* genomes annotation 9
Phylogenomic analyses 10
Analysis of symbiont contribution to amino acid biosynthesis pathways 10
Microscopic analyses 11
Histological and ultrastructural analyses 11
Fluorescence *in situ* hybridization 11
**Supplementary Text** **11**
**Challenges and unusual observations in fast-evolving symbiont genomes annotation** 11
Very low similarity between homologous regions 11
Gene truncation in *Sulcia* genomes 12
Open reading frames of distinct genes merged into one ORF 13
Open reading frames broken up into putative separate genes 14

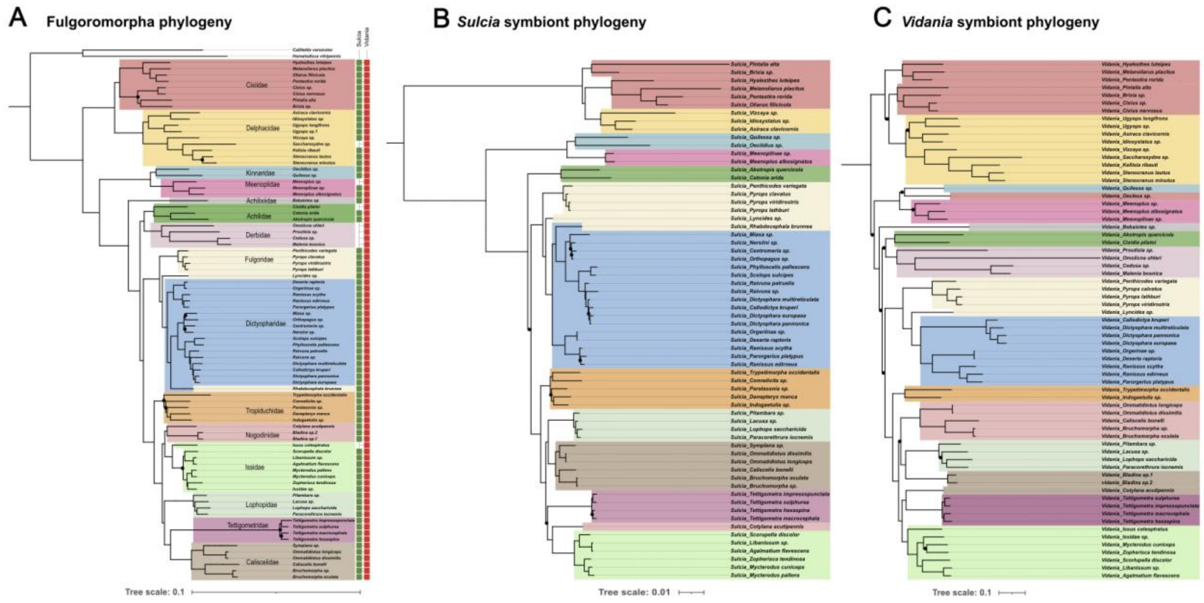


**Figure S1. Co-diversification of planthoppers and their ancestral endosymbionts *Sulcia* and *Vidania*, based on genome-level data.** The phylogenies of ancient bacterial endosymbionts *Sulcia* and *Vidania* essentially recapitulate that of their planthopper hosts, indicating strict co-diversification. **A.** The maximum likelihood phylogeny of 149 planthopper species based on 1164 nuclear and 13 mitochondrial genes (Deng et al. 2024); the topology is based on all species regardless of the infection state, but only the specimens that host either *Vidania* or *Sulcia* are shown; **B.** The ML phylogeny of 68 *Vidania* strains based on 98 genes; **C.** *Sulcia* ML phylogeny based on 130 genes. In all phylogenies, we used the same set of colors to indicate planthopper families. Black circles on the branches indicate bootstrap support values below 100.

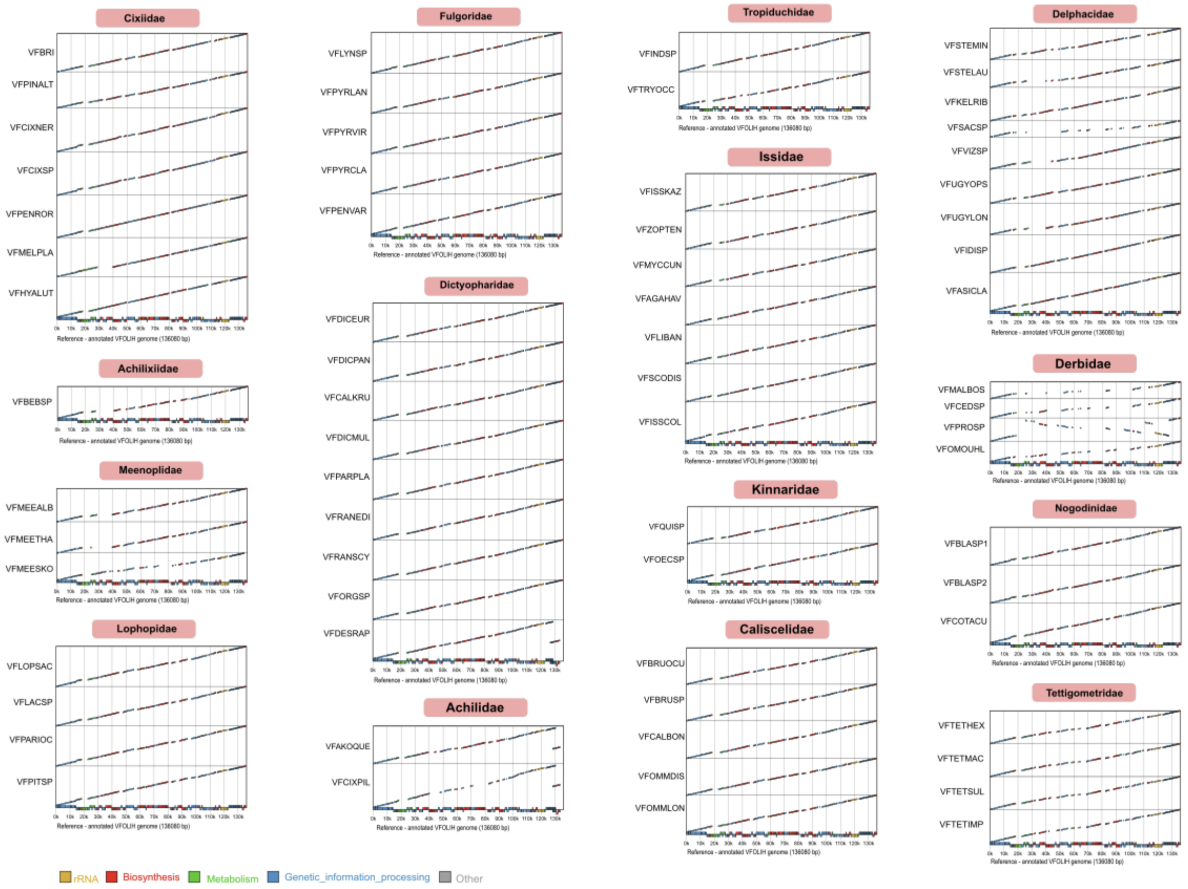


**Figure S2.** The alignment of all completed *Vidania* genomes against one of the most complete in the collection - that of *Vidania* VFOLIH from *Oliarus filicicola* (Cixiidae) - shows that the genomes generally retain conserved organization (with one exception - strain VFPROSP, Derbidae), but substantial portions of the genome are often lost. Alignment plots were made using a custom workflow that used annotation information.

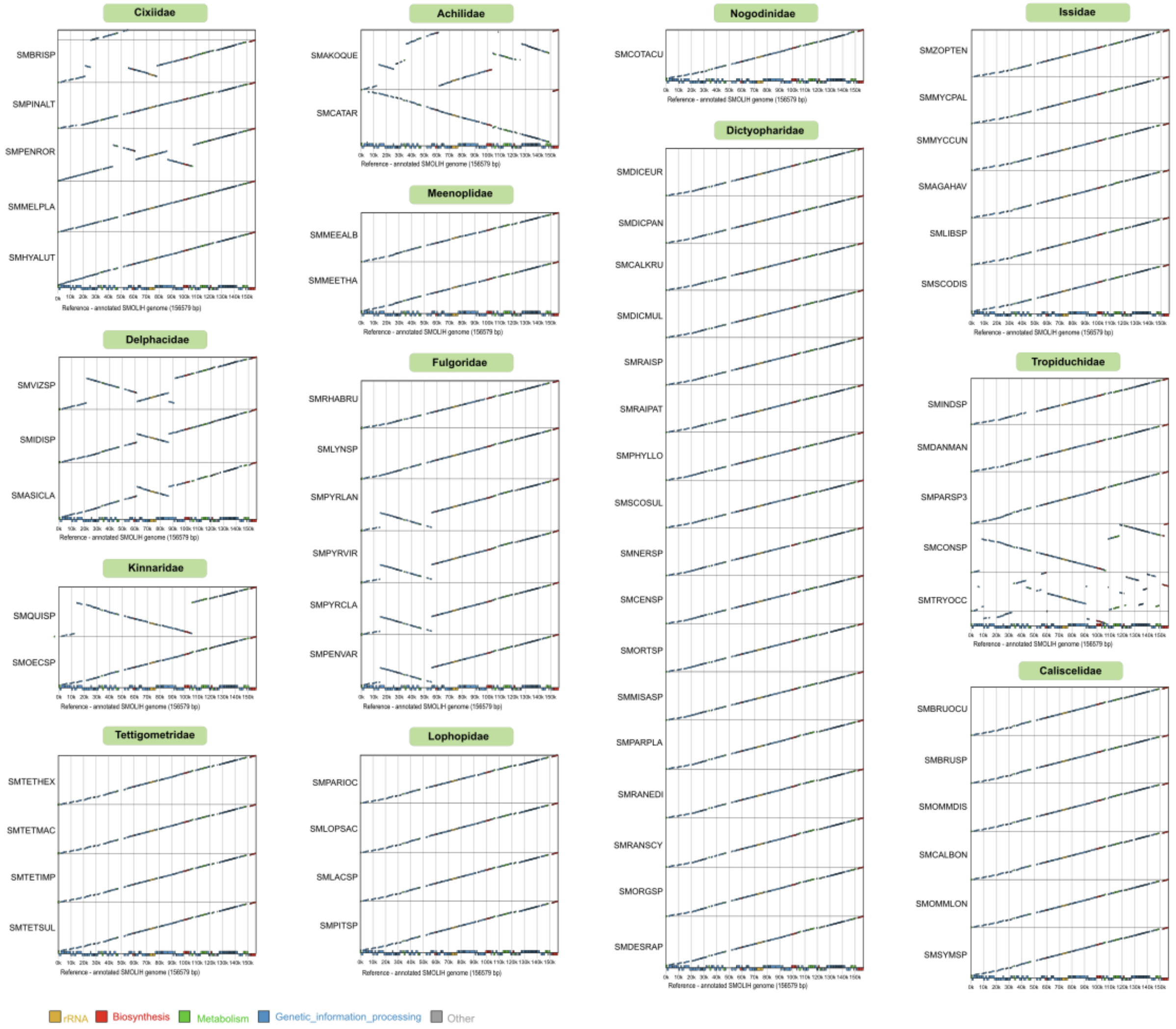


**Figure S3.** The alignment of all completed *Sulcia* genomes against one of the most complete in the collection - that of *Sulcia* SMOLIH from *Oliarus filicicola* (Cixiidae) - shows that the genomes usually retain conserved organization, but there have been multiple independent cases of genomic rearrangements, sometimes multiple. Alignment plots were made using a custom workflow that used annotation information.

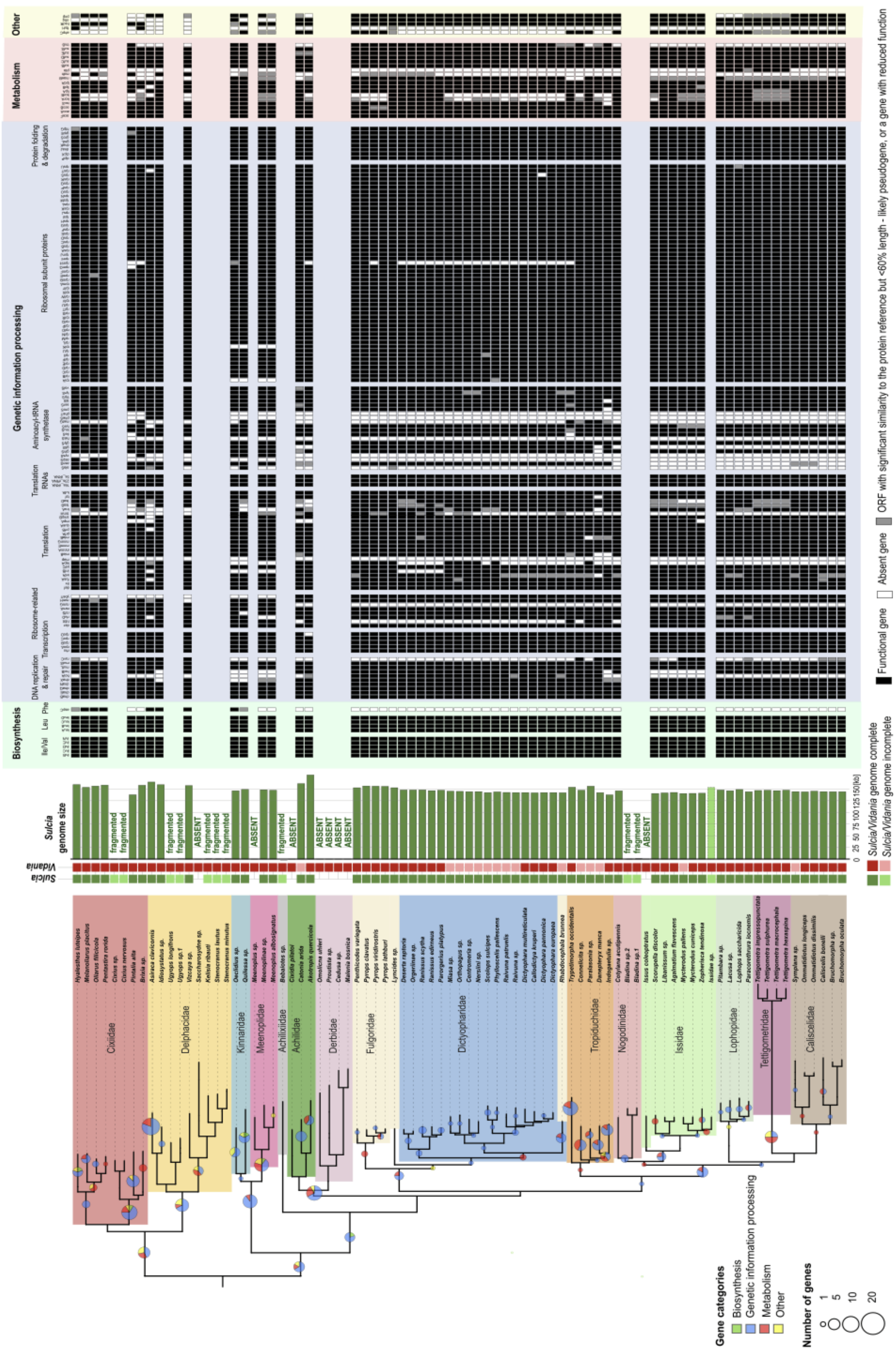


**Figure S4.** The distribution of genes across all complete symbiont *Sulcia* genomes from different planthopper species. As in Fig. 4, the insect phylogeny was redrawn from Fig. 1 but trimmed to retain only the clades where either *Sulcia* or *Vidania* genome is complete. For all complete *Sulcia* genomes and all genes with known functions, we show whether the gene was identified, and either deemed complete or truncated by >40% relative to the reference alignment (putative pseudogene). Circles on tree branches indicate the number of genes that were reconstructed as lost on a particular branch.

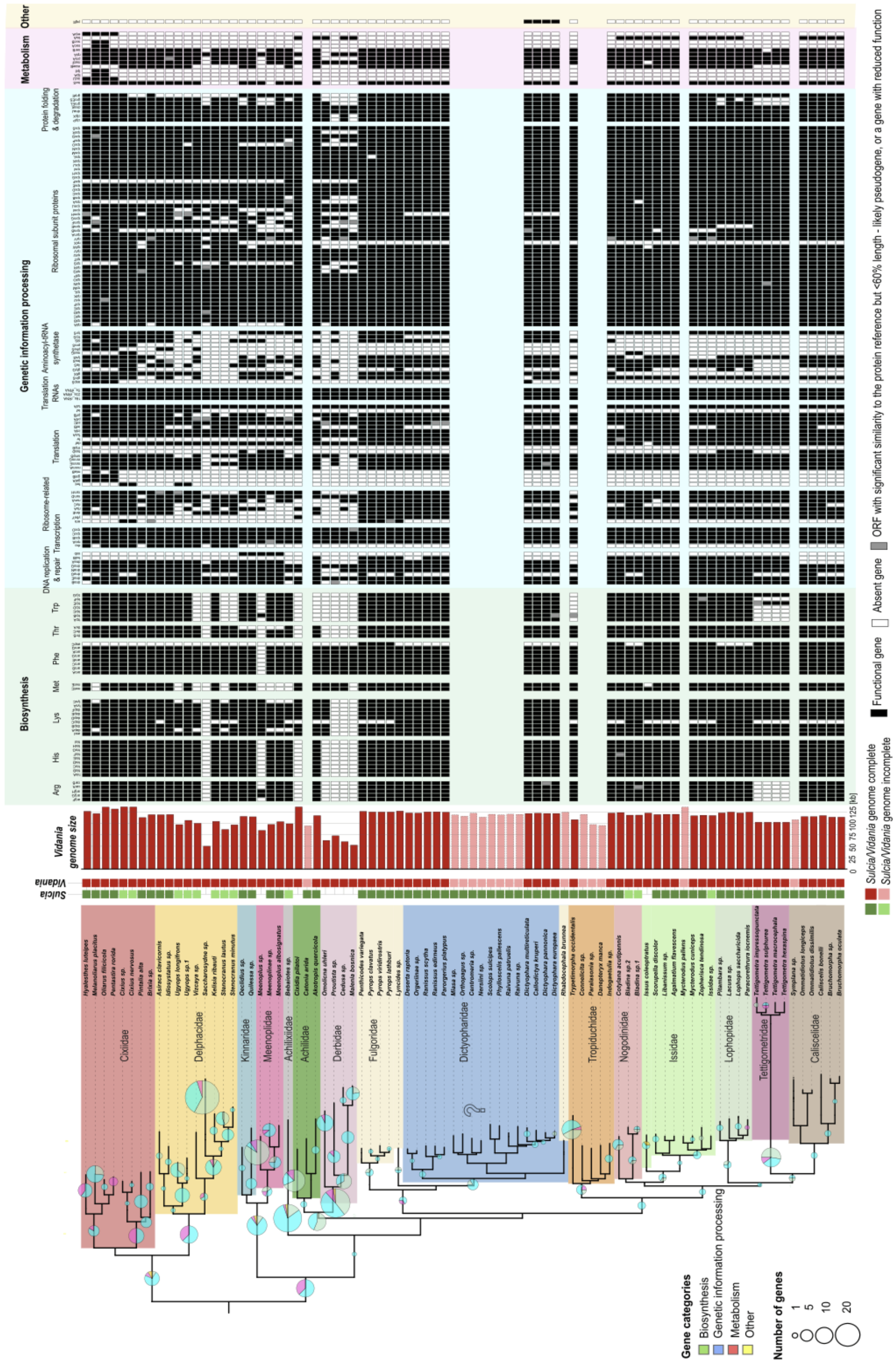


**Figure S5.** The distribution of genes across all complete symbiont *Vidania* genomes from different planthopper species. As in Fig. 4, the insect phylogeny was redrawn from Fig. 1 but trimmed to retain only the clades where either *Sulcia* or *Vidania* genome is complete. For all complete *Vidania* genomes, and all genes with known functions, we show whether the gene was identified, and either deemed complete, or truncated by >40% relative to the reference alignment (putative pseudogene). Circles on tree branches indicate the number of genes that were reconstructed as lost on a particular branch.

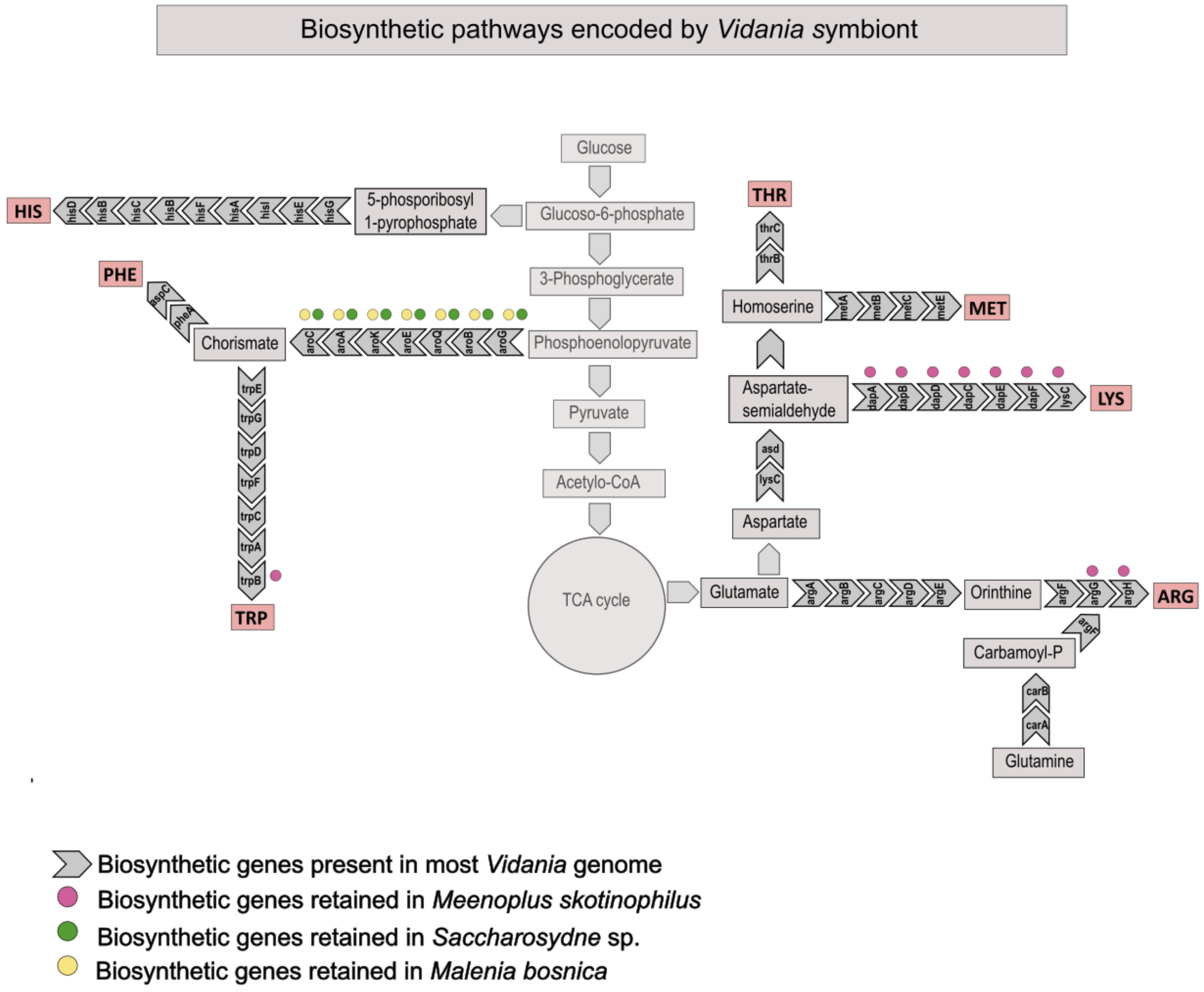


**Figure S6.** The representation of the ancestral amino acid biosynthetic pathways encoded by the ancestral *Vidania* - and the subsets encoded by the smallest *Vidania* genomes from three recognized superfamilies (VFSACSP1 from *Saccharosydne* sp., Delphacidae; VFMALBOS from *Malenia bosnica*, Derbidae; VFMEESKO from *Meenoplus skotinophilus*, Meenoplidae).

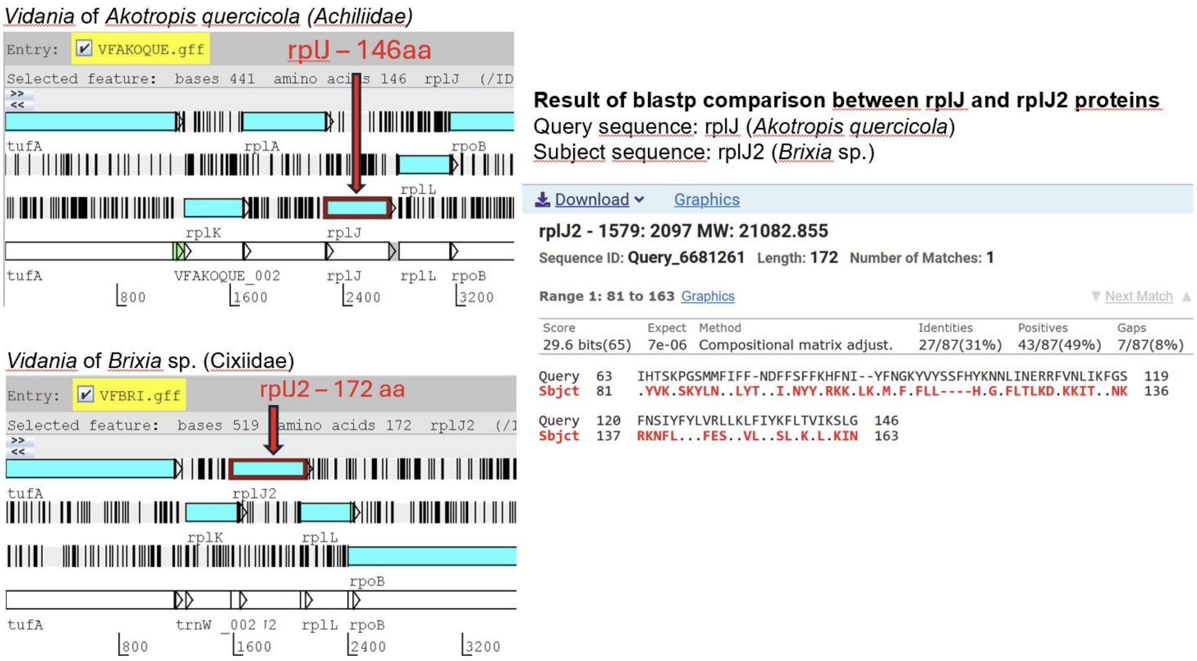


### Gene truncation in *Sulcia* genomes

In *Sulcia*, certain genes were often substantially truncated relative to the most common length in the dataset, while retaining high sequence similarity. The truncation was often phylogenetically conserved, seen in all species from some clades, suggesting selective pressure for the retention of truncated variants and thus their functionality. We conclude that during the process of genome reduction, some gene domains may have been lost, but the truncated protein retained enough of the original function to justify their long-term retention. For such truncation conserved in at least 3 species, we established alternative reference alignments to ensure that they are scored as “functional”.

**Example:** The size of rplI gene (ribosomal protein) in *Sulcia* genomes from families Kinnaridae, Fulgoridae, Dictyopharidae, and Tropiduchidae is around 450 nucleotides, very similar to its length in *Eschericha coli* (447 nucleotides - https://www.uniprot.org/uniprotkb/P0A7R1/entry) whereas in the remaining nine families, we observed the truncation of this gene down to around 200 nucleotides.

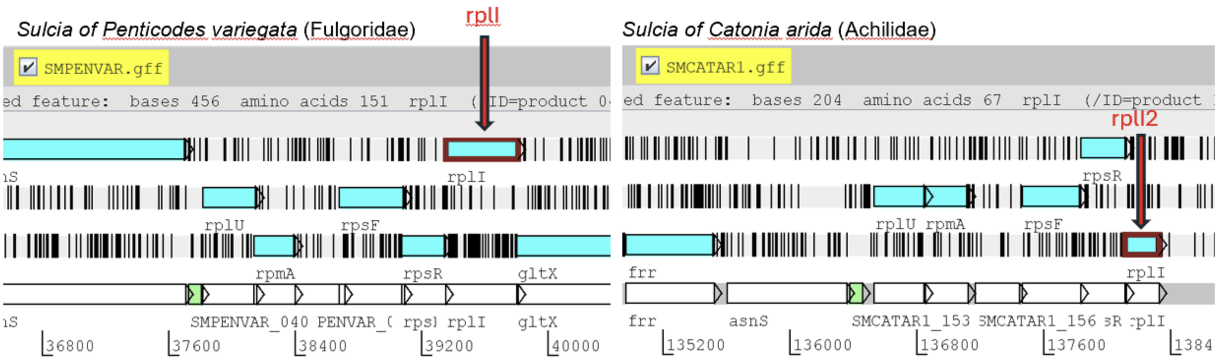


### Open reading frames of distinct genes merged into one ORF

In *Vidania*, we observed cases when adjacent ORFs representing different genes, with very different functions, became merged into a single ORF. These cases were typically phylogenetically conserved - observed in symbionts in all species from a clade. Considering that the merged ORF contained nearly full-length coding sequences of the two genes, we concluded that both genes are likely functional and result in two distinct functional proteins. To ensure their successful detection, we created HMM references for the merged ORFs.

**Example:** In *Vidania* genomes from the clade comprising families Caliscelidae, Nogodinidae, Tettigometridae, Lophopidae, and Issidae, ORFs of genes tilS (tRNA(Ile)-lysidine synthase - a genetic information processing gene) and lysC (aspartate kinase - an amino acid biosynthesis gene), adjacent but separate in other planthoppers, became merged into a single large ORF. When such an ORF was detected, we scored both genes as present.

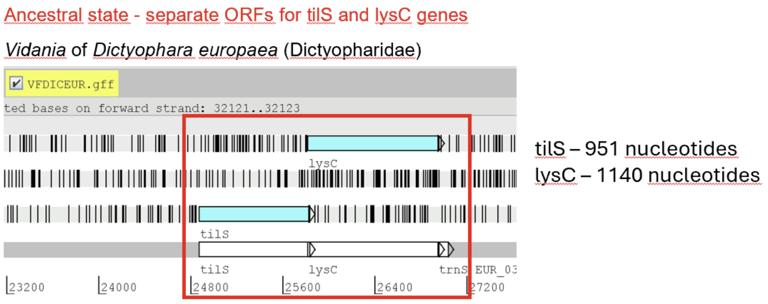


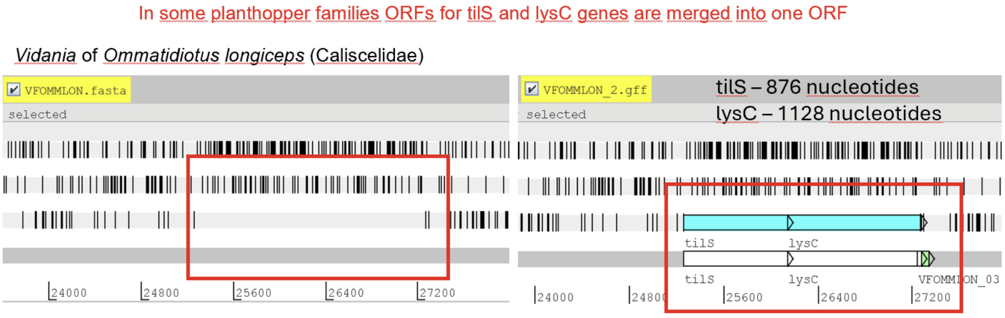


### Other examples:

- in *Vidania* genomes from the family Lophopidae, adjacent genes *dapC* and *dapD* were merged
- in *Vidania* genomes from the family Cixiidae, adjacent genes *maeB* and *aroE* were merged.

### Open reading frames broken up into putative separate genes

In several genomes, we observed STOP codons or frameshifts within the open reading frames of conserved genes, confirmed as the dominant variant by inspecting mapped reads. In several cases, we excluded sequencing or assembly errors. We expect that in some cases, this might represent a recent mutation that has spread but not reached fixation within the highly polyploid symbiont genome, or within a population of symbiont cells. We also hypothesize that sequences may sometimes be corrected post-transcriptionally through an unknown mechanism. However, when considering such genes, we used the standard criteria, where ORFs of at least 60% of the reference length were scored as functional, and the shorter ones were scored as "truncated - putative pseudogenes".

**Example:** Gene *dnaE* (DNA polymerase III alpha subunit) in *Vidania* of *Zopherisca tendinosa* is broken by an apparent frameshift into two separate ORFs, the longer of which was ca. 72% of the reference length. In this case, we classified the gene as functional.

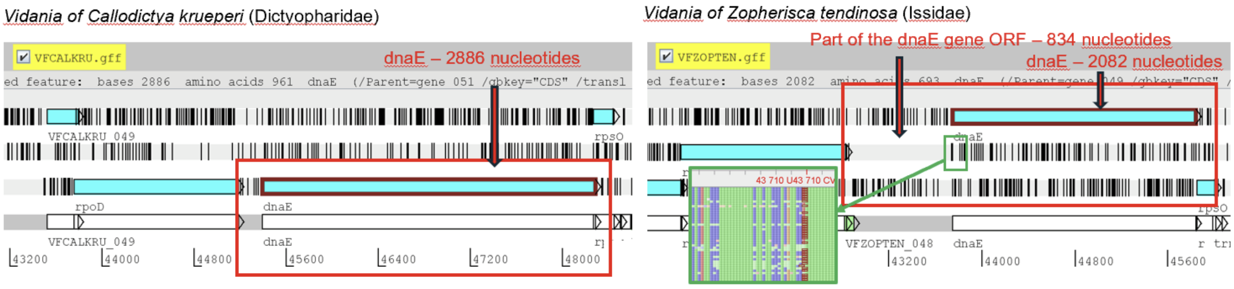
